# Alveolar-Basal Intermediates Drive Pulmonary Fibrosis via Coordination of a Pro-Fibrotic Signaling Niche in Silicosis

**DOI:** 10.64898/2026.03.11.711170

**Authors:** Barbara Zhao, Helen I. Warheit-Niemi, Kathleen C. S. Cook, Lori Pitstick, Andrea Toth, Amber Elitz, Francis X McCormack, Amanda L. Zacharias, William J. Zacharias

## Abstract

Pulmonary fibrosis is a progressive, terminal disease with high morbidity. Existing therapeutics slow disease progression but do not reverse fibrotic lung remodeling, accentuating the importance of identifying cellular mechanisms that underlie lung fibrosis. Recent literature suggests that alveolar type 2 (AT2) progenitors undergo transition to stressed Krt8^high^ cells following lung injury. Accumulation of stressed Krt8^high^ cells is a hallmark of acute and chronic lung diseases, particularly pulmonary fibrosis. Whether Krt8^high^ cells actively participate in the fibrotic process or are simply an epiphenomenon of lung injury remains unclear. We previously described a genetic model in which deletion of the lung transcription factor Nkx2-1 in the AT2 progenitor lineage induces AT2 progenitors to transition to the Krt8^high^ cell state. Here, we use this tractable genetic model to directly test the pathogenic influence of Krt8^high^ cells. We show that Nkx2-1^-/-^ Krt8^high^ cells accumulate in a Krt7^high^/Krt19^high^/Krt17^neg^ alveolar-basal intermediate cell state (ABI). Following induction of fibrotic lung injury with inhaled silica, ABI enter a unique inflammatory state (iABI) that drives fibrotic remodeling via coordination of a fibrotic signaling niche containing inflammatory alveolar fibroblasts (iAF) and pulmonary osteoclast-like cells (POLC). Computational analysis predicts that iABI elaborate pro-inflammatory signals which increase matrix deposition by iAF and induce differentiation of interstitial macrophages to a profibrotic POLC variant. Niche mapping demonstrates that iABI, iAF, and POLCs interact within newly formed fibrotic niches in the lung alveolus in mice with pre-existing accumulation of ABI. These data provide direct evidence that ABI accumulation in fibrotic lung disease is pathogenic.

## Introduction

The two major epithelial cell types present in the alveolar epithelium of all mammals are alveolar epithelial type 1 (AT1) and alveolar epithelial type 2 (AT2) cells. AT1 cells contact capillary endothelial cells to make up the gas exchange surface of the lung, while AT2 cells produce pulmonary surfactant to prevent atelectasis during ventilation^1^. During acute lung injury, the respiratory barrier is compromised; AT2 cell proliferation and subsequent AT2 to AT1 cell trans-differentiation are required to re-epithelialize the alveolus to restore barrier function and gas exchange capacity^2-5^. AT2 progenitors, also called alveolar epithelial progenitors (AEPs), are mobilized by Wnt and Fgf signaling following lung injury to undergo expansion and differentiation and regenerate the alveolar epithelium^6-9^. The process of AT2 to AT1 differentiation is critical to lung regeneration, leading to a recent expansion of literature assessing AT1 differentiation following infectious, toxic, and genetic lung injuries. These studies have characterized an intermediate cell state during AT2 to AT1 cell differentiation which exhibit hallmarks of cell stress, arise following bleomycin lung injury, IL1b-induced inflammation, AT1-specific ablation via diphtheria toxin, and hyperoxia during influenza A infection^10-15^. While named variously by multiple groups, common to all descriptions of this cell state is elevated expression of Keratin 8 (Krt8), a cytoskeletal marker which is broadly expressed at low levels across all lung epithelia and more highly expressed in proximal epithelial cells and upregulated in lung adenocarcinoma^16,17^. Krt8 is more highly expressed in these stressed transitional epithelial cells, so we denote them here as Krt8^high^ cells. The heterogeneity of Krt8^high^ cells is unclear, though extensive evidence from human disease samples has implicated similar cell states in acute and chronic lung diseases^18-21^. Given the robust induction of Krt8^high^ cells following diverse lung injuries and their accumulation in human lung disease, there is a strong rationale to better understand the mechanisms causing Krt8^high^ cell accumulation and their role in the diseased lung.

Krt8^high^ cells were originally identified and have been extensively evaluated in the context of fibrotic lung injury. Recent data has demonstrated that Krt8^high^ cells upregulate integrin β6 (Itgb6) and contribute to mechanisms of fibrosis via activation of myofibroblasts^22^ or increased TGFβ expression^15,23,24^. Krt8^high^ cells encompass an apparently heterogeneous group of transitional cell states with names that include the pre-alveolar type-1 transitional cell state (PATS), alveolar-basal intermediate (ABI) 1 and ABI2 cell states^25^. ABI, particularly ABI2 cells, are enriched in severely fibrotic regions of IPF lungs compared to control lungs and non-fibrotic regions of IPF lung samples^10-12,26-28^. Existing studies have evaluated the induction and molecular function of Krt8^high^ cells during alveolar stress in the context of murine lung injuries^10-12^. However, these lung injury models cause damage to not only the alveolar epithelium but also existing mesenchymal, endothelial, and immune cells in the lung. Krt8^high^ cell interactions with fibroblasts and immune cell types are likely central to the role of Krt8^high^ cells in lung disease, but are challenging to directly assess during injuries that simultaneously damage all of these compartments in the lung. Additionally, broad immune activation in the context of extensive lung injury makes it difficult to assess whether epithelial stress drives a worsening inflammatory milieu or vice versa. These challenges have limited the current evidence, which has to date not addressed whether Krt8^high^ cell accumulation prior to injury, as in patients with established lung disease, is protective or deleterious as the lung responds to injury and regenerates.

In this work, we directly test the role of accumulated Krt8^high^ cells on the progression of lung fibrosis. We previously described a tractable model for studying Krt8^high^ cell accumulation, in which genetic ablation of Nkx2-1 in AEPs (Nkx2-1^KO^) promotes irreversible acquisition of a Krt8^high^ state during lung homeostasis^13^. This model allowed us to generate mice in which Krt8^high^ cells are present prior to lung injury, enabling direct evaluation of the impact of accumulated Krt8^high^ cells during injury and regeneration on the alveolar niche. Here, we show that Nkx2-1^KO^ Krt8^high^ cells accumulate in a state most similar to previously described ABI/ABI1 cell state characterized by a combination of Krt8^hi^/Krt7^hi^/Krt19^hi^ expression. To isolate the effect of pre-existing ABI during injury, we chose to induce fibrosis with inhalation of crystalline silica, which causes fibrotic lung injury in both murine models^29-31^ and human patients^32,33^. Chronic inhalation of silica particulate in human patients manifests as a spectrum of lung disease including progressive massive fibrosis, diffuse interstitial fibrosis, or simple nodular fibrosis^34^. The pathophysiology of silicosis in human patients is driven primarily by chronic injury to alveolar macrophages, where RANK ligand and inflammatory cytokine activation drive macrophage differentiation towards pulmonary osteoclast-like cells (POLCs) which potentiate fibrosis. This pathophysiology is recapitulated in murine models of silicosis 28-56 days after silica administration^35^.

Here, we show that silica-induced fibrosis occurs without spontaneous induction of Krt8^high^ cells in control animals, allowing direct assessment of the impact of accumulated ABI on lung fibrosis in Nkx2-1^KO^ mice. Following silica-induced fibrotic lung injury, Nkx2-1 KO ABI take on a unique inflamed ABI (iABI) state, which then alters the alveolar microenvironment. iABI cell activation potentiates a physical and signaling niche containing inflammatory alveolar fibroblasts (iAF), activated alveolar monocytes, and POLCs. Within this fibrotic niche, iABIs engage in crosstalk with both iAFs and osteoclastic macrophage populations to amplify pro-fibrotic and inflammatory signaling. These data directly implicate ABI accumulation and persistence in a fibrotic niche driving lung disease.

## Results

### Krt8^high^ cell accumulation prior to silicosis increases lung fibrosis severity

To evaluate the epithelial-driven influence of Krt8^high^ cells in the fibrotic lung milieu, we first generated Krt8^high^ cells *in vivo* via tamoxifen-induced recombination using the AT2 progenitor-enriched *Tfcp2l1*^*CreERT2*^ driver in combination with *Nkx2-1*^*flox/flox*^, as previously described^13^. *In vivo* genetic ablation of *Nkx2-1* in AT2 progenitors leads to acquisition of a stressed cell state characterized by high Krt8 expression and loss of SFTPC expression in lineage-labelled cells within 2 weeks of tamoxifen treatment^13^. To induce fibrotic lung injury, we administered silica particles intratracheally 2-, 4-, or 8 weeks after tamoxifen induction of Krt8^high^ cell and harvested the lungs for downstream analyses 28 days after silica administration [**Fig.1A**]. Silica treatment in both wild-type litermates (WT) and Nkx2-1^KO^ mice caused significant weight loss that peaked 2 days post-injury (dpi), with recovery of weight in both groups by 28dpi [**Supplementary Fig.1A**].

**Figure 1.**
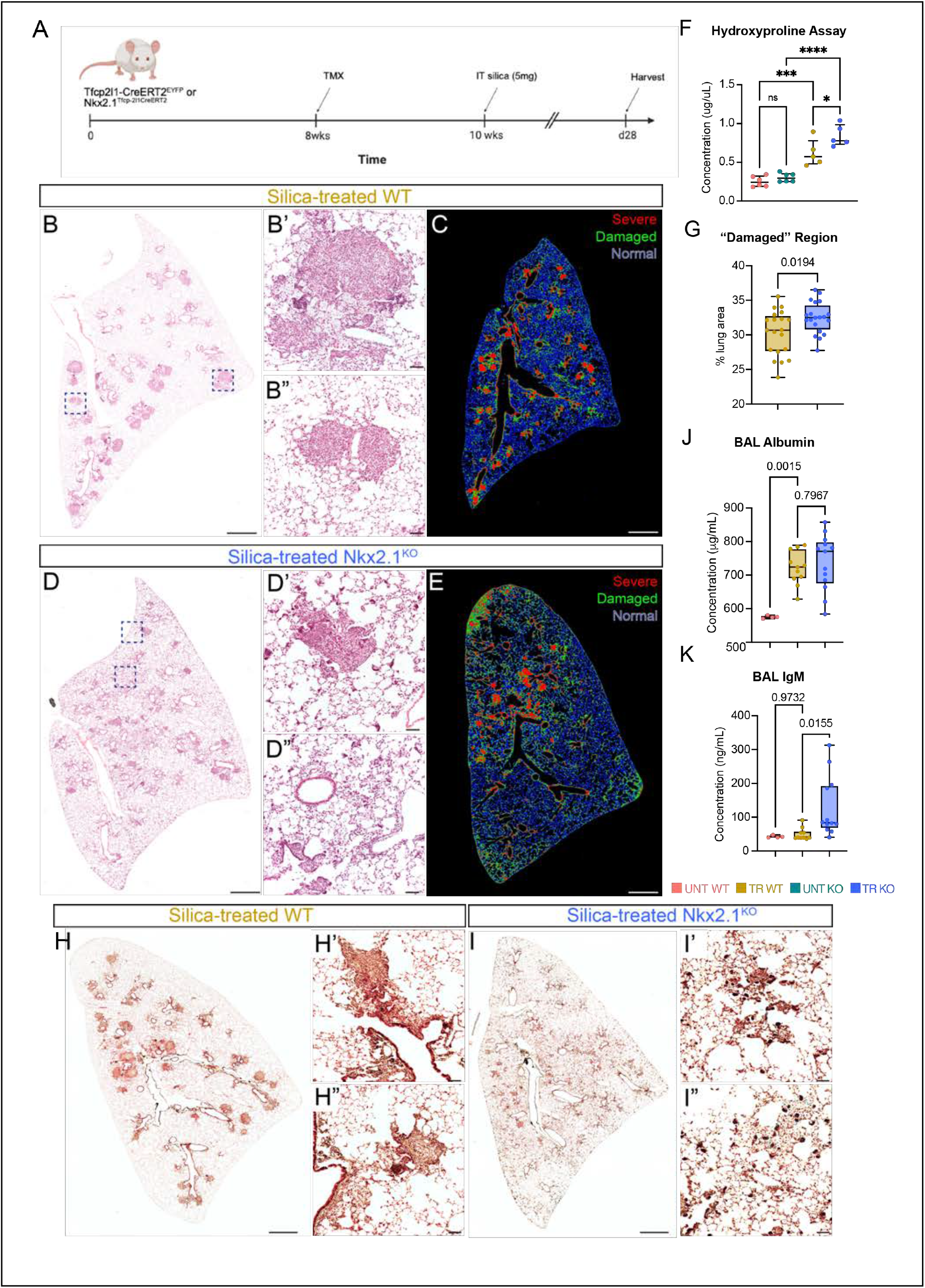
Pre-injury, accumulated Krt8^high^ epithelia are associated with increased lung damage, respiratory barrier disruption, and lung fibrosis in experimental silicosis. (A) Experimental schematic. (B-E) H&E staining demonstrates silica-induced lung injury areas are more discrete in silica-treated (TR) WT (B) and more widespread in TR Nkx2-1 KO (D). Lung damage assessment program showed increased “damaged” lung surface area (green) in TR KO lungs (E) compared to TR WT lungs (C); this is quantified in (G) with exact p values calculated by t-test. (F) The hydroxyproline assay revealed significantly increased collagen deposition in TR animals compared to untreated (UNT) animals, and significantly increased collagen in TR KO compared to TR WT. Exact p-values calculated by one-way ANOVA with multiple comparisons. (H-I) Pentachrome staining of TR WT lungs (H) and TR Nkx2-1 KO lungs (I) demonstrates increased collagen and elastin deposition in TR KO lungs compared to TR WT lungs. Collagen and reticular fibers (yellow), elastic fibers (black), mucin (blue), and fibrin (red). (J) Albumin concentration in the bronchoalveolar lavage fluid (BALF) was significantly increased in both TR WT and TR KO animals compared to UNT animals, but no differences were observed in albumin concentration between TR WT vs TR KO animals. (K) IgM concentration in the BALF was significantly increased in TR KO compared to TR WT animals. Exact p-values calculated by one-way ANOVA with multiple comparisons.

We observed increased lung damage in Nkx2-1^KO^ lungs at 28dpi compared to WT mice exposed to silica. Histological examination by H&E showed widespread fibrotic remodeling more expansive than the typical silicosis nodules found in WT animals [**Fig.1B-E**], which correlated with significantly increased hydroxyproline content in Nkx2-1^KO^ animals [**Fig.1F**]. Quantification of lung damage using clustering of image pixels to characterize distinct zones of lung injury as “severe,” “damaged,” and “normal” in an unbiased manner^36^ showed that Nkx2-1^KO^ lungs displayed a significantly higher percentages of “damaged” lung surface area compared to silica-treated WT lungs [**Fig.1B-1E**,**G**]. Percentages of “severe” lung injury between the two groups were not different; severe injury was typically scored in areas of fibrotic nodules [**Supplementary Fig.1B**]. We observed increased collagen deposition in Nkx2-1^KO^ lungs compared to control lungs with pentachrome staining [**Fig.1H-I**]. We then assessed alveolar epithelial barrier disruption in Nkx2-1^KO^ by quantifying IgM and albumin concentrations in murine bronchoalveolar lavage fluid (BALF)^37-40^. Neither albumin, 70kDa^37,41^, nor IgM, 900kDa, are detected in the alveolar airspace during homeostasis [**Fig.1J-K**] and their presence in the BALF therefore indicates increased permeability of the respiratory barrier. Albumin concentrations were significantly increased in BALF from both silica-treated WT and Nkx2-1^KO^ mice compared to untreated mice, indicating alveolar epithelial barrier disruption in silica-induced lung injury [**Fig.1J**]. Interestingly, IgM concentrations were not significantly increased in silica-treated WT BALF compared to untreated mice, but were markedly increased in silica-treated Nkx2-1^KO^ BALF compared to untreated mice of both genotypes and silica-treated WT animals [**Fig.1K**]. These data demonstrated that silica injury disrupts the respiratory barrier and that silica injury in the presence of Krt8^high^ cells leads to further barrier disruption allowing IgM to leak into the alveolar space. Taken together, these data demonstrated that accumulated Krt8^high^ cells exacerbated fibrosis and impaired barrier function following silica-induced lung injury.

We hypothesized that increased severity of lung injury with accumulated Krt8^high^ cells was attributable to the differential impact of Krt8^high^ cells in the alveolar niche. We therefore evaluated the localization and cell states in the Nkx2-1^KO^ Krt8^high^ cell lineage following silicosis. Lineage-labelled Nkx2-1^KO^ cells were Krt8^hi^/SFTPC^low^ following silicosis, and we noted large clusters of these cells preferentially located in areas of silica-induced lung injury in Nkx2-1^KO^ mice [**Fig.2A-2D**] especially in alveolar regions distinct from fibrotic nodules. Assessment of lineage-labelled AEPs in WT animals following silica treatment showed areas of cell shape change and proliferation surrounding areas of severe fibrosis. However, these lineage-labeled cells were Krt8^low^/SFTPC^hi^, with few Krt8^high^ cells near fibrotic areas. [**Fig.2A-2D**].

**Figure 2.**
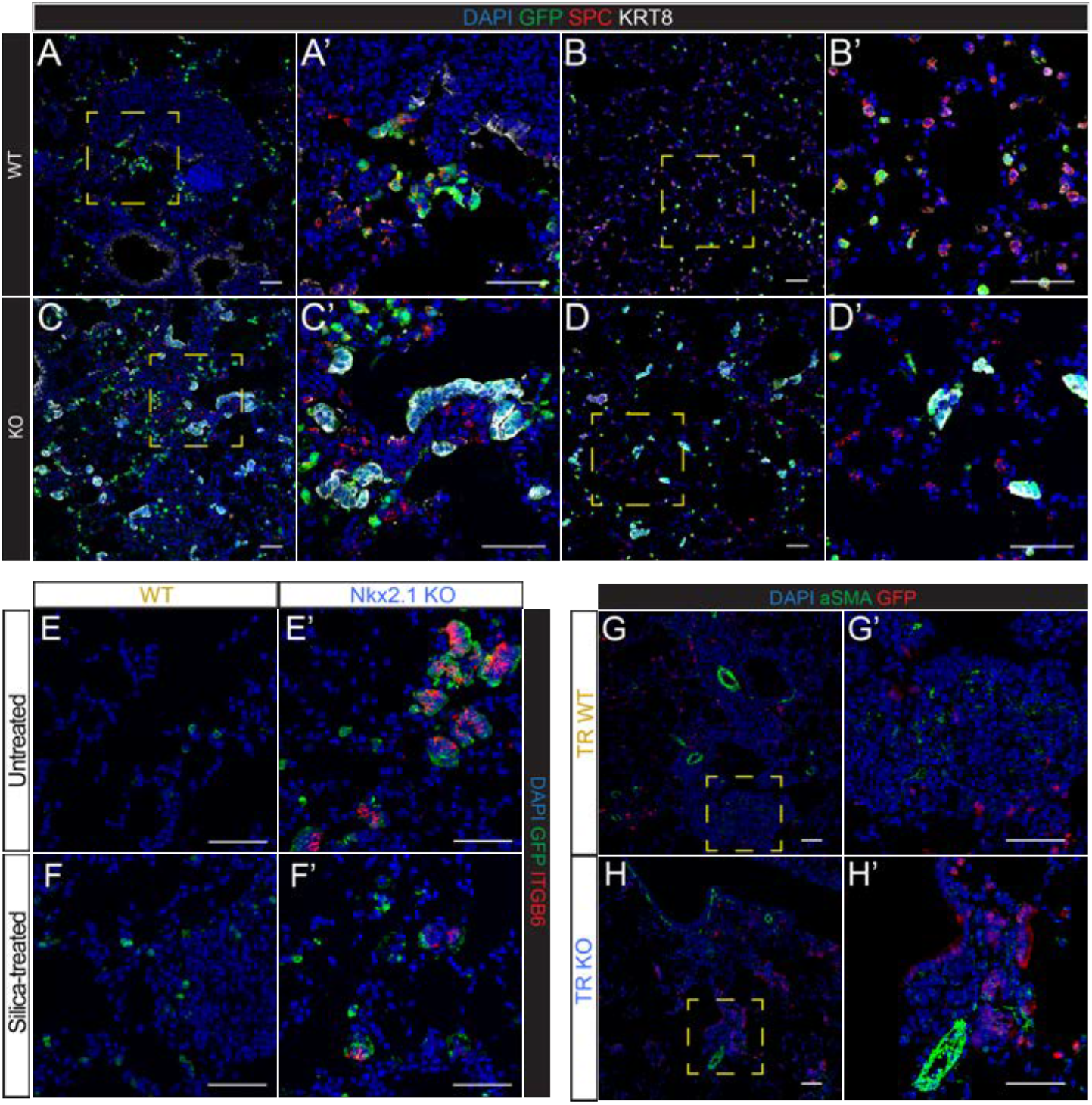
Krt8^high^ epithelia clusters near areas of silica-induced lung injury. (A-D) Lineage-labeled GFP*+*/Krt8^high^/SPC*-* epithelial clusters preferentially surround areas of lung injury in TR KO lungs (C), whereas lineage-labeled GFP*+*/Krt8^low^/SPC*+* AEPs are found adjacent to areas of lung injury in TR WT (A). AEPs and Krt8*hi* epithelial clusters are located in uninjured areas of TR lungs (B,D). (E-F) Integrin b6 (Itgb6) expression is selectively observed in Krt8^high^ cell clusters that are present in UNT and TR KO lungs. aSMA staining was observed in injured areas of both TR WT (G) and TR KO (H) lungs.

Krt8^high^ cells have been reported to express multiple cell junctional markers, such as E-cadherin, Claudin4 (Cldn4), and integrin β6 (Itgb6), suggesting differential cell-cell interaction potential in the Krt8^high^ population compared to AT2 progenitors^10,11,13^. Nkx2-1 KO epithelial cell clusters express high levels of ITGB6, E-cadherin and CLDN4^13^, prior to silica treatment which persisted throughout the course of silicosis [**Fig.2E-F**]. This expression pattern is consistent with the hypothesis that Nkx2-1 KO lineage cells alter cell-cell interactions in the fibrotic niche, as Itgb6 expression in epithelium has been shown to correlate with activation of Tgfβ signaling, and loss of Itgb6 expression is protective against the development of pulmonary fibrosis^23^. Concordantly, we observed close association of lineage labeled Krt8^high^ cells with alpha-smooth muscle actin (aSMA) expression in injured areas of silica-treated Nkx2-1 KO lungs, indicating activation of myofibroblasts after silica injury [**Fig.1G-1H**]. Interestingly, comparison of silicosis severity 2, 4, or 8 weeks following tamoxifen showed no significant difference in severity of injury or degree of fibrosis [**Supplementary Fig.2A-H**], suggesting that presence of Krt8^high^ cells, rather than length of exposure of the lung to Krt8^high^ cells, was the primary factor conferring increased severity after silica response. Together, these data are consistent with the model that Nkx2-1 KO-lineage Krt8^high^ cells directly drive fibrotic niche organization in silicosis.

### Krt8^high^ cells in the Nkx2-1 KO lineage are alveolar-basal intermediates (ABI)

Multiple studies indicate substantial heterogeneity among Krt8^high^ cells following lung injury. To characterize the Krt8^high^ cell states observed in our murine Nkx2-1 KO model, we used fluorescence-activated cell sorting (FACS) to isolate lineage-labelled Nkx2-1 null Krt8^high^ cells from untreated Nkx2-1 KO lungs and lineage-labelled AT2 progenitors from WT lungs and conducted single-cell RNA sequencing (scRNA seq) to compare the molecular state of Nkx2-1 null Krt8^high^ cells [**Fig.3A-B**].

**Figure 3.**
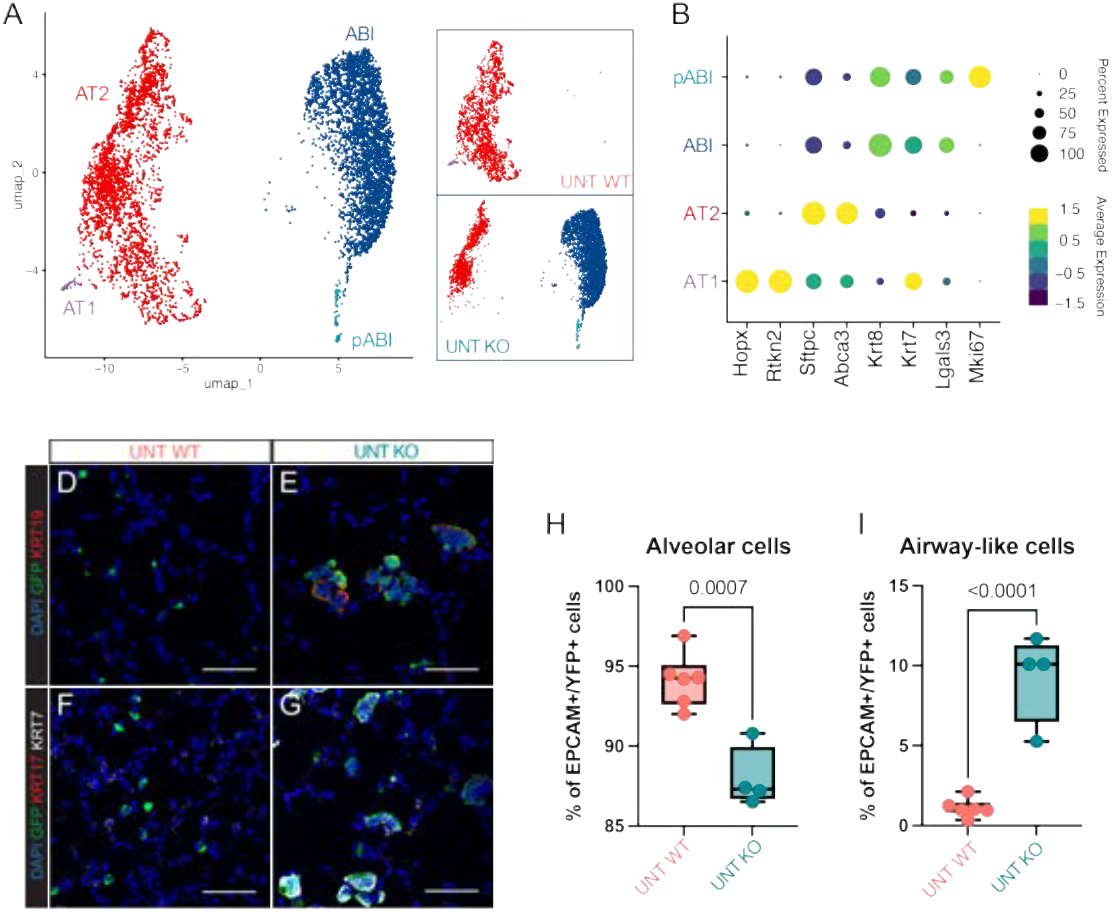
Krt8^high^ epithelia are in an alveolar-basal intermediate (ABI) cell state. (A) UMAP identifying epithelial cell states from untreated WT and Nkx2-1 KO conditions. (B) Dotplot expression markers for AT1, AT2, ABI, and proliferating ABI (pABI) epithelial states. (C-F) Untreated Nkx2-1 KO clusters are Krt7^high^/Krt19^high^/Krt17^neg^ compared to untreated WT AEPs. Flow cytometry shows that alveolar cell fate acquisition is significantly decreased in UNT KO lungs compared to UNT WT (G) and that airway cell fate is correspondingly significantly increased (H).

Integrated analysis of FACS-sorted lineage-labeled epithelial cells from untreated WT and Nkx2-1 KO mice identified 4 distinct epithelial cell states through Seurat graph-based clustering [**Fig.3A**]. As expected, FACS-sorted AEPs in WT animals were predominantly AT2 cells defined by canonical expression of AT2-associated genes (i.e., *Abca3, Lamp3*, and *Sftpc*) [**Fig.3A-B**], with a small AT1 cell state defined by canonical expression of AT1-associated genes (i.e., *Hopx* and *Rtkn2*). The Nkx2-1 KO lineage contained a small proportion of AT2 cells and no detectable AT1s. The lineage otherwise consisted of Krt8^high^ cells, which were present exclusively in Nkx2-1 KO animals. These cells showed minimal *Sftpc* expression and significantly increased expression of *Krt7, Krt8*, and *Krt19* [**Fig.3A-B**]. These findings were confirmed via immunofluorescence (IF), which showed expression KRT7, KRT8, and KRT19, and lack of KRT17 expression [**Fig.3D-3G**] in Nkx2-1 KO lineage labeled epithelial cells. Flow cytometry for alveolar (Epcam^hi^/CD31^low^/YFP^+^/Itgb4^low^/CD24^low^) and airway (Itgb4^high^/CD24^high^) cell states [**Supplementary Fig.3**] demonstrated that many cells in the Nkx2-1 KO lineage had adopted airway-like cell surface marker expression [**Fig.3H-I**]. However, Nkx2-1 null cells rarely differentiated into ciliated, secretory, or basal cells *in vivo* [**Supplementary Figure 4A-I]**. Taken together, these data demonstrate the Nkx2-1 KO lineage Krt8^high^ cells are in a molecular state most consistent with previously described alveolar-basal intermediate cells (ABI).

### Nkx2-1 KO ABI cells undergo inflammatory activation in the silica-induced fibrotic lung milieu

We next evaluated the evolution of the AT2 lineage in both WT and Nkx2-1 KO animals following silica-induced lung injury. Integrated analysis of FACS-sorted AEPs or ABIs from untreated WT and untreated Nkx2-1 KO, as well as silica-treated WT and silica-treated Nkx2-1 KO mice, identified 5 distinct epithelial cell states through Seurat graph-based clustering [**Fig.4A**]. We noted a newly identified cell state which arose exclusively in the silica-treated Nkx2-1 KO condition [**Fig.4A-4B, 4E**], with increased expression of inflammatory response genes suggesting an activated inflammatory state; we denoted this state as inflamed ABI (iABI). We hypothesized that the iABI state was arising secondary to lung injury.

**Figure 4.**
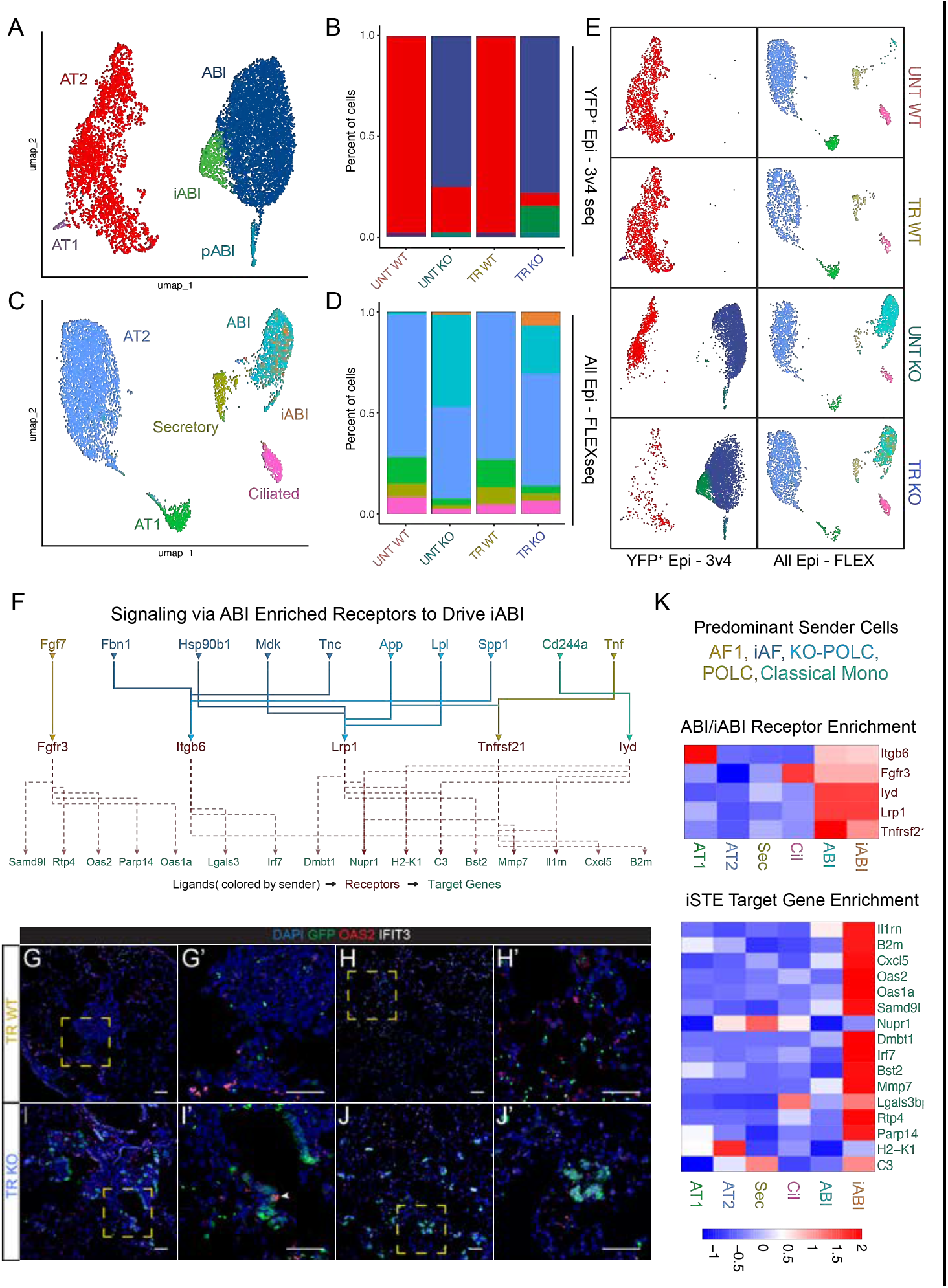
An inflamed ABI (iABI) cell state emerges after silica treatment. (A) UMAP of integrated 3v4 scRNA sequencing analysis of FACS-sorted lineage-labeled epithelial cells from untreated (UNT) WT, UNT Nkx2-1 KO, silica-treated (TR) WT, and TR Nkx2-1 KO conditions identified 5 distinct epithelial cell states. (B) Frequency of epithelial cell state in each condition. Inflamed ABIs (iABI) are exclusively present in TR Nkx2-1 KO condition. (C) UMAP of epithelial subset of whole lung scRNA sequencing via 10X Flex Genomics from all conditions identifies 6 distinct epithelial cell states. (D) iABIs and ABIs are present only in KO epithelial subsets. (E) UMAPs of integrated scRNA sequencing analysis of FACS-sorted lineage-labeled epithelia and epithelial subset from whole lung scRNA sequencing separated by condition; ABIs are only present in Nkx2-1 KO conditions and iABIs are only present in silica-treated Nkx2-1 KO condition in both data sets. (F) NicheNet analysis of whole lung scRNA sequencing reveals ligands produced by alveolar fibroblasts (AF), inflammatory AF (iAF), pulmonary osteoclast-like cells (POLC), and classical monocytes predicted to act on ABI-enriched receptors (red, K) to drive expression of iABI markers of inflammation (green, K). (G-J) OAS2 and IFIT3 expression is observed in GFP*+* ABIs in TR KO condition in injured areas (arrows) consistent with iABI state; iABIs are not observed in uninjured areas of TR WT and TR KO. (K) Enriched receptors and gene expression markers of ABIs and iABIs compared to other identified epithelial cell states.

To further evaluate the driving signals leading to iABI cell state acquisition and the cellular interactions of iABIs within the fibrotic alveolar niche, we conducted whole lung single-cell RNA sequencing with 10X Genomics Chromium Single Cell Gene Expression Flex on all experimental conditions [**Supplementary Fig. 5**]. Integrated analysis of the epithelial cell subset identified all the distinct epithelial cell states present from FACS-sorted scRNA-sequencing, as well as additional ciliated and secretory airway epithelial cell states [**Fig. 4C-4E**]. Combined analysis of FACS-sorted YFP^+^ cells with 3v4 RNA sequencing and whole lung FLEX data confirmed that ABI were present exclusively in Nkx2-1 KO conditions and that iABI were present exclusively in the silica-treated Nkx2-1 KO condition [**Fig.4E**]. To define upstream signals driving iABI acquisition, we evaluated ligand-receptor interactions between ABIs and other cell states in the alveolar niche using a combination of CellChat^42,43^ and NicheNet^44^. ABI receive signals from multiple cell types within the fibrotic niche (i.e., *Fbn* and *Mdk* from alveolar fibroblasts, *App* and *Spp1* from POLCs) that activate ABI-enriched receptors (Itgb6, Lrp1), and these ligand-receptor interactions drive expression of iABI markers associated with inflammation [**Fig.4F**]. Inflamed ABIs retained canonical ABI markers including *Krt7/Krt19* and were defined by expression of inflammatory markers, such as *Oas2, Ifit3, Irf7, Ifi44*, and *Isg15. Lgals3*, a key marker of Krt8^high^ cells following bleomycin injury, was also highly upregulated in iABI [**Fig.4K**] Further analysis of genes enriched in iABI cells demonstrated broad expression of genes associated with immunomodulatory GO terms including response to interferon-beta and the regulation of innate immune response **[Supplemental Figure 6]**. We identified ABI cells within fibrotic lung regions by IF for the iABI enriched targets OAS2 and IFIT3, and identified multiple iABI in GFP^+^ lineage-labelled Nkx2-1 KO cells [**Fig.4G-4J**]. ABI and iABIs were notably absent in control animals [**Fig.4G-4J**], supporting the conclusion that signaling driven by iABI cells may underlie the severe fibrosis seen in Nkx2-1 KO animals.

### iABI cells drive inflammatory alveolar fibroblast (iAF) activation during silicosis

We next sought to evaluate the consequences of iABI presence in fibrotic niches in the Nkx2-1 KO lung. Among epithelial cells, iABIs specifically expressed a number of pro-fibrotic and pro-inflammatory ligands [**Fig.4F,K**], including *Rankl, Ccn1, Spp1, Mmp7*, and *Cxcl5*. We therefore reasoned that these signals may participate in sculpting the milieu of the lung during silicosis-induced fibrosis. We first evaluated the mesenchymal compartment, were recent data has demonstrated that alveolar fibroblasts differentiate towards inflammatory (iAF) fibroblast state marked by expression of LCN2 and SAA3 followed by a fibrotic fibroblast (fAF) state marked by expression of CTHRC1^45^. Consistent with prior reports in silicosis^45^, we identified very few *Cthrc1*-expressing iAF cells by scRNAseq. Integrated analysis of the mesenchymal subset of whole lung scRNA sequencing data from all experimental conditions instead identified seven distinct mesenchymal cell states [**Fig.5A**], with a prominent component of inflammatory alveolar fibroblasts. These iAF were defined by expression of *Meox2, Saa3, Lcn2*, and *Ccl2*, and emerged in both WT and Nkx2-1 KO lungs after silica treatment [**Fig.5B-5C**]. iAF frequency was higher in silica-treated Nkx2-1 KO lungs compared to silica-treated WT lungs [**Fig.5B**], suggesting that their emergence may be influenced by the presence of ABI/iABI. We identified iAF in both WT and Nkx2-1 KO lungs after silica-induced lung injury by IF. iAF co-expressed SAA3, LCN2, and MEOX2 [**Fig.5D-5G**], and were enriched in the most severe fibrotic lesions in animals from both genotypes. Gene expression of *Lcn2* was highest iAFs from silica-treated Nkx2-1 KO mice [**Fig.5H**]. We then evaluated ligand-receptor interactions that drive iAF emergence and function in the fibrotic niche. We found that signaling ligands expressed on ABIs, iABIs, and myeloid lineages including POLCs were predicted to interact with iAF-enriched receptors to activate an iAF-enriched gene signature which included a pro-fibrotic program driven by *Osteopontin/Spp1* and an inflammatory response program driven by TNF and IL1 signaling. Together, these inputs are predicted to increase expression of key iAF target genes that contribute to ECM deposition and immune cell recruitment [**Fig.5I**]. Furthermore, analysis of genes enriched in iAFs relative to AF1 and AF2 cells identified GO terms and KEGG pathways relating to the inflammatory response, positive regulation of cytokine production and ECM organization **[Supplemental Figure 7]**. iABI-enriched ligands, which included *Spp1, Ccn1*, and *Mmp7* were predicted to contribute to ECM deposition downstream of iAF, consistent with a known role of these pathways in pulmonary fibrosis in both humans^46-50^. Many of the ligands expressed by iAF were predicted to signal to pulmonary macrophages, so we next turned our attention to the macrophage and monocyte compartment of the iABI/iAF niche.

**Figure 5.**
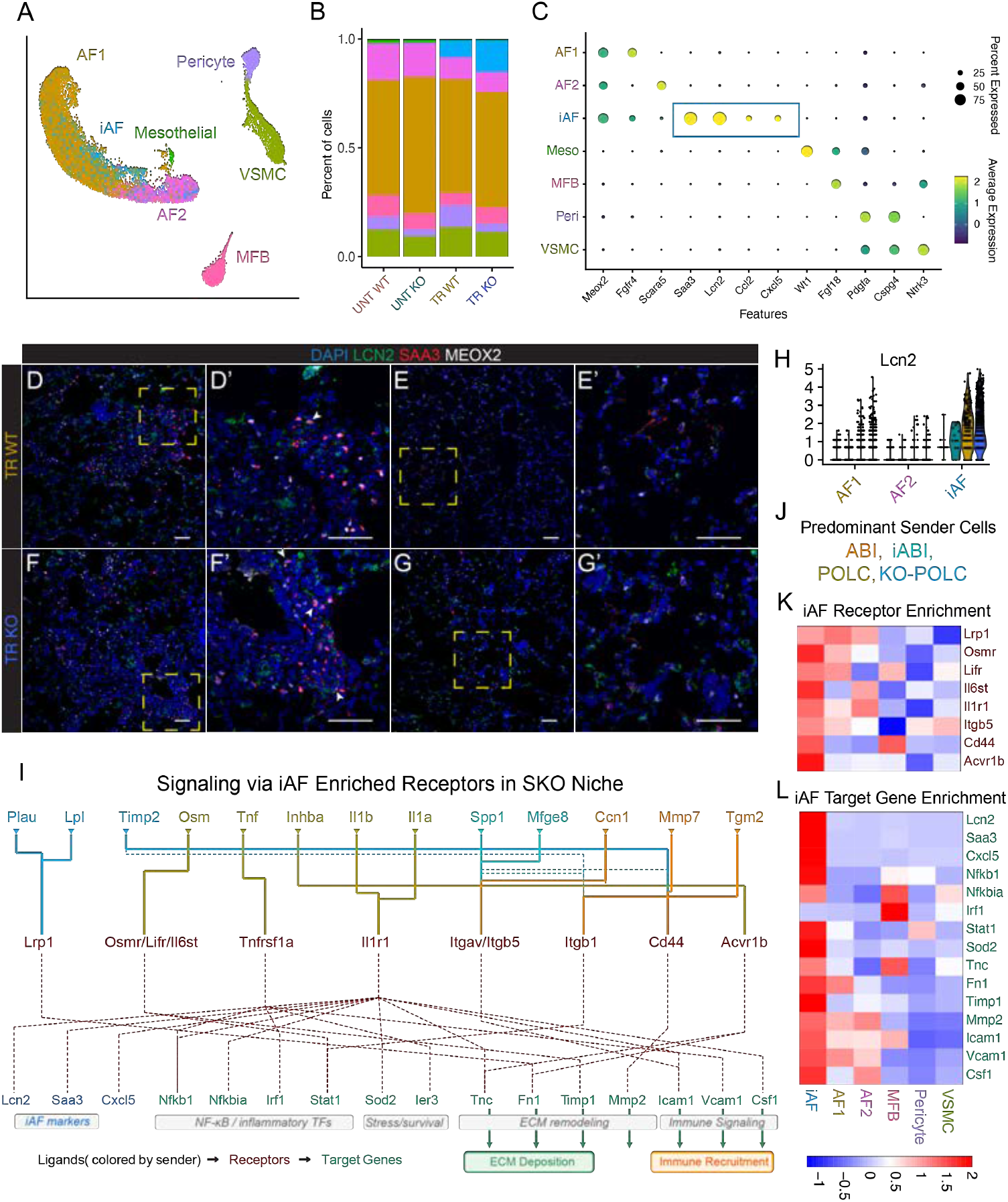
Inflammatory alveolar fibroblasts (iAF) emerge after silica-induced lung injury. (A) UMAP of mesenchymal subset from whole lung sequencing data identifying 7 distinct mesenchymal cell states. (B) Mesenchymal cell frequencies demonstrate increased number of iAF1 in TR Nkx2-1 KO lungs compared to TR WT lungs. (C) Dotplot shows gene expression markers for each mesenchymal cell state with upregulation of inflammatory markers in iAFs. (D-G) Immunohistochemistry demonstrates iAF presence (arrows) with co-expression of SAA3, LCN2, and MEOX2 exclusively in injured areas of TR WT and TR KO lungs. (H) Violin plot of *Lcn2* mRNA in iAFs in UNT WT, UNT KO, TR WT, and TR KO shows highest expression of *Lcn2* in TR KO. (I) NicheNet analysis demonstrates that ligands produced by ABI, iABI, and POLCs (J) act directly on iAF-enriched receptors (K) which regulate genes involved in iAF identity, ECM deposition, and immune cell recruitment (L).

### ABI drive differentiation of pro-fibrotic POLC from interstitial macrophages to potentiate pulmonary fibrosis

Macrophages play a prominent role in the regulation of lung fibrosis, as monocyte-derived alveolar macrophages are recruited to regions of alveolar injury and have been demonstrated to contribute to fibrotic tissue remodeling^51-53^. In silicosis, both alveolar macrophages and interstitial macrophages are known to transition to an osteoclast-like (POLC) state driven by RANKL/RANK signaling which contribute to development of fibrosis following silica exposure in both mice and humans. The pathogenic role of POLCs is further supported by the finding that reducing POLC differentiation via blockade of RANKL improves fibrosis following silica exposure^35^. Integrated analysis of immune cell populations from our whole lung sequencing data revealed increased numbers of alveolar macrophages (AM), interstitial macrophages (IM), and POLCs [**Fig.6A-6B**]. Consistent with previous data^35^, multiple macrophage subsets demonstrated upregulation of a POLC-like gene program, with two populations identifiable as POLCs [**Fig.6C]**. One of these populations, denoted POLCs, was present in both WT and Nkx2-1 KO mice following silicosis, while a second, denoted KO POLC, was present almost exclusively in Nkx2-1 KO.

**Figure 6.**
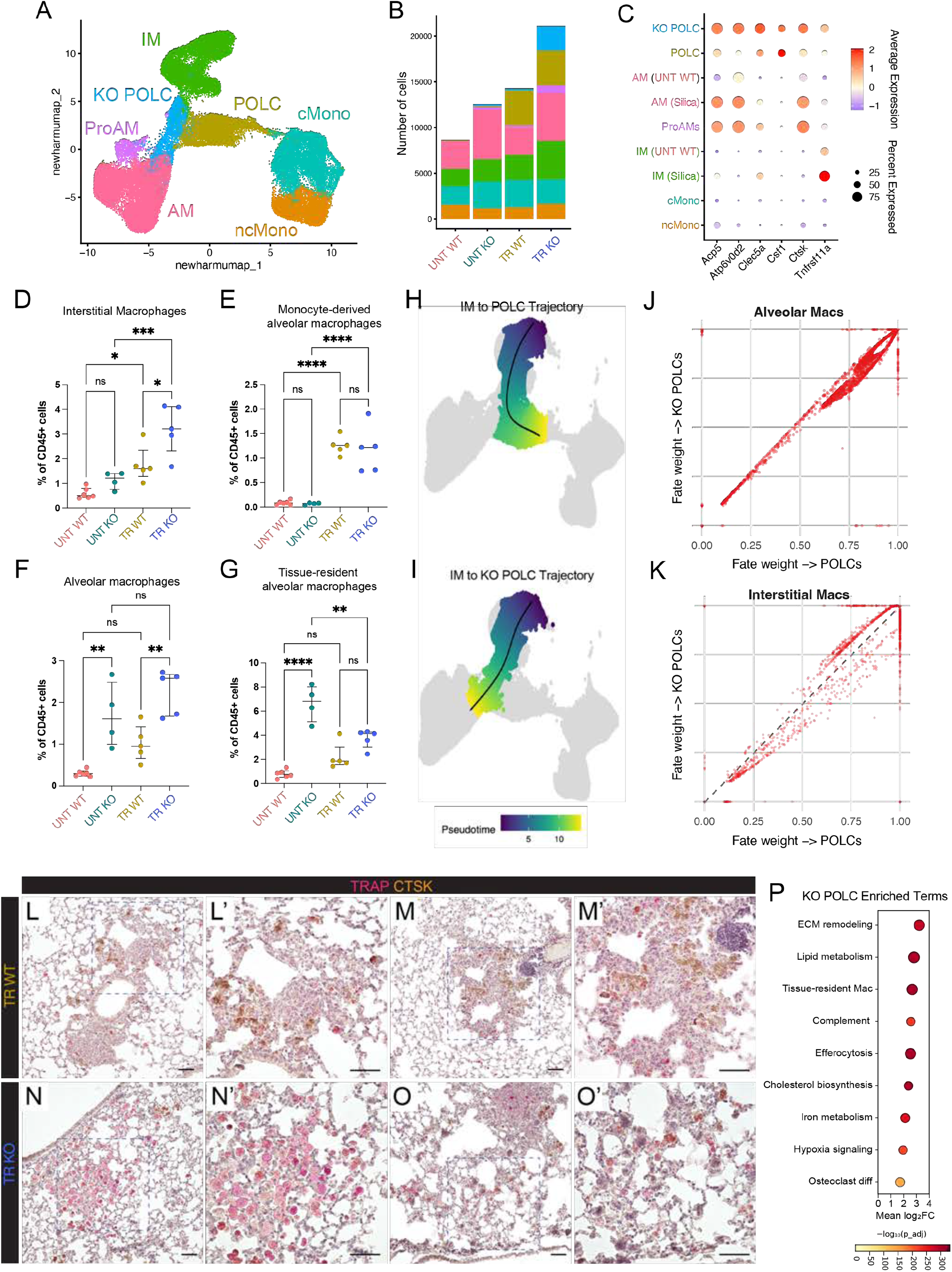
ABI drive differentiation of interstitial macrophages toward pro-fibrotic pulmonary osteoclast-like cells (POLCs) to potentiate fibrosis following silica. (A) UMAP detailing macrophage/monocyte cell populations from integrated analysis of whole lung scRNA sequencing concatenated across all experimental conditions. (B) Cell population frequencies show upregulated alveolar macrophage (AM), interstitial macrophage (IM), POLC, and KO-specific POLC frequencies TR Nkx2-1 KO condition. (C) Dotplot of osteoclast-like gene expression markers show upregulated expression of osteoclast-like genes in silica-associated AMs, IMs, and POLCs, with further upregulated expression in conditional KO-specific POLCs. (D-G) Flow cytometry delineates upregulated AM frequencies as tissue-resident as opposed to monocyte-derived; tissue-resident interstitial macrophages (IM) are significantly increased in TR KO compared with TR WT lungs. (H-I) Pseudotime analysis shows that POLCs and KO POLCs predominantly emerge from IM, which are upregulated in the TR KO condition. (J-K) Fate probability maps show POLCs arise from both IM and AM, while KO POLCs are likely derived from IMs. (L-O) immunohistochemistry demonstrating that TRAP*+*/CTSK*+* POLCs and KO POLCs are enriched throughout injured areas of TR lungs. KO POLCs exhibit higher expression of TRAP and with POLC-like morphology (N-O). (P) Gene Ontology (GO) terms enriched in KO POLCs compared to other POLCs from all conditions demonstrate enrichment of genes involved in pro-fibrotic processes like ECM remodeling, inflammation, and osteoclast differentiation.

We confirmed changes in multiple macrophage subsets using spectral flow cytometry^54^ following silica-induced lung injury [**Fig. 6D-G**]. We identified small but significant increases in tissue-resident interstitial macrophages (IM) in both silica-treated WT and silica-treated Nkx2-1 KO conditions compared to untreated conditions [**Fig. 6D**]. We also observed large increases in AM frequencies in both untreated and silica-treated Nkx2-1 KO lungs [**Fig.6E-G**]; these AMs were largely CD64^hi^/CD11b^low^ tissue-resident AMs rather than monocyte-derived AMs [**Fig.6E-6G, Supplementary Figure 8**]. Thus, although exposure to silica does increase monocyte-derived AMs, the increase in tissue-resident AMs is much larger in the silica treated Nkx2-1 KO condition.

To evaluate the origin of the POLC and KO POLC cell states, we performed trajectory analysis with Slingshot^55^ [**Fig.6H-I**] and condition-specific trajectory topology evaluation [**Fig.6J-K**] with Condiments^56^. Both AM and IM populations were predicted as possible cells of origin for POLCs in both WT and Nkx2-1 KO mice following silica administration [**Fig.6J-K**]. Conversely, KO POLCs were predicted to arise almost entirely from IM rather than AM [**Fig.6J-K**], consistent with the increased numbers of IMs seen by flow cytometry and scRNA-seq. Supporting this conclusion, KO POLCs show expression high expression of key markers genes for interstitial macrophages such as *C1qa* and *C1qb*, while POLCs do not [**Supplemental Figure 9]**. We therefore predicted that the ABI/iABI niche would be enriched for POLCs. To that end, we performed Tartrate-Resistant Acid Phosphatase (TRAP) and cathepsin K (CTSK) staining to localize POLCs in the lung [**Fig.6L-O**]. We detected POLCs in fibrotic nodules in both WT and Nkx2-1 KO mice. We also noted large numbers of POLCs in areas of peripheral fibrotic remodeling throughout the Nkx2-1 KO lung, while few POLCs were present in the WT lung outside of fibrotic nodules [**Fig.6L-O**]. Comparison of gene expression in KO POLC compared to POLC showed significant enrichment of genes associated with fibrosis, hypoxemia, and inflammatory signaling [**Fig.6P, Supplemental Figure 9]**, further suggesting that KO POLCs in the Nkx2-1 KO niche potentiated fibrosis.

### A feed forward, pro-fibrotic loop coordinated by ABIs following silicosis

In support of this conclusion, combined CellChat and NicheNet analysis suggested that POLCs, KO POLCs, and iABIs were among the most strongly enriched signaling centers in the overall signaling milieu of the Nkx2-1 KO lung following silicosis [**Fig.7A**]. Multiple pro-fibrotic pathways were notably enriched in this milieu, including RANKL, Osteopontin/SPP1, TNF, and TGFβ [**Fig.7B**]. POLCs were poised to respond to many of these pro-fibrotic signals [**Fig.7C**], with prominent inputs from iABI and iAF implicated in activation the pro-fibrotic and pro-inflammatory programs which defined the KO POLCs state. These silica-specific, ABI-driven KO POLCs produced multiple ligands (Spp1, App, Rankl, Csf1) that likely signal to iABI and iAF to stimulate pro-inflammatory and pro-fibrotic milieu [**Fig.7C, Supplemental Figure 10**].

**Figure 7.**
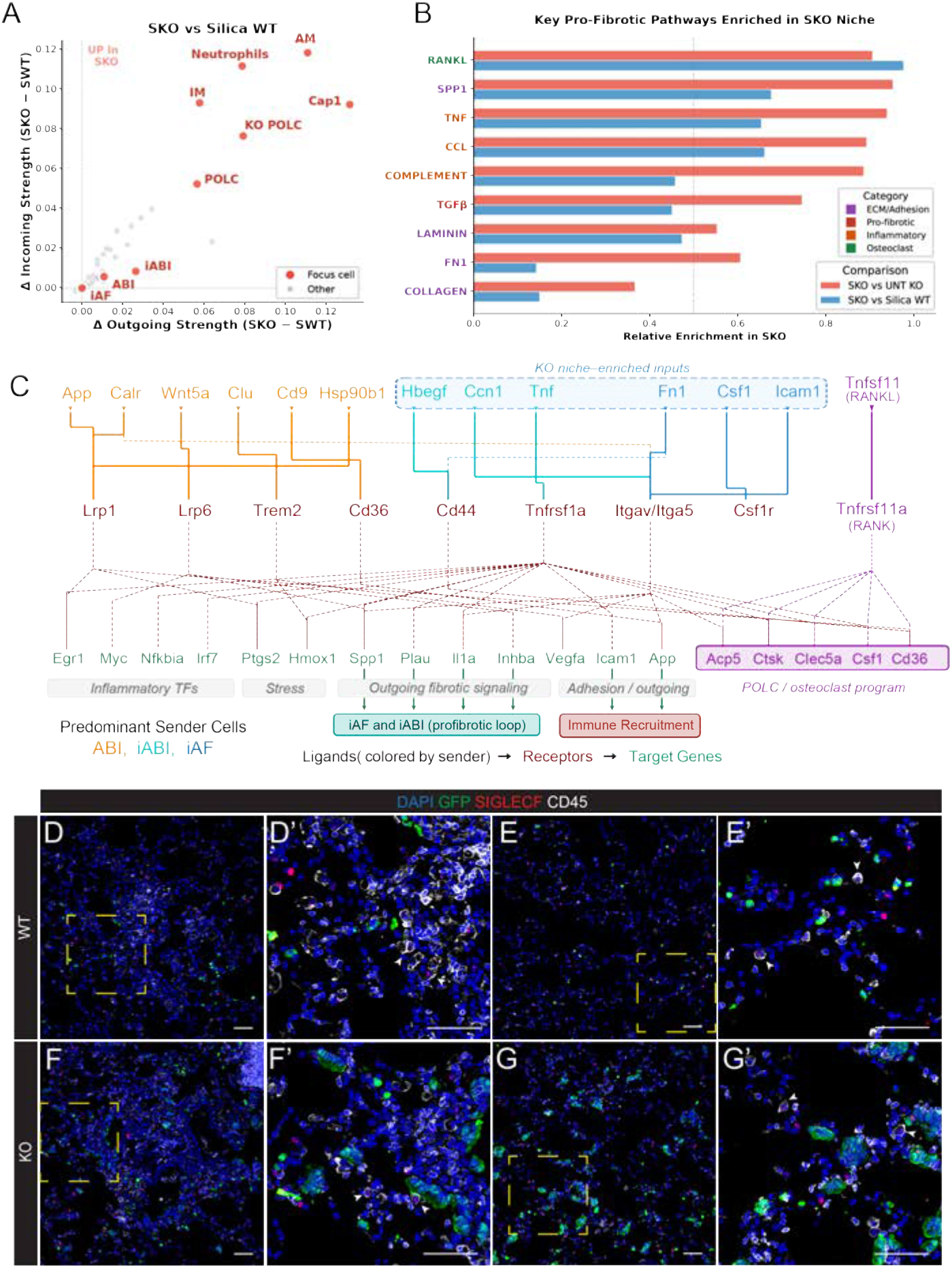
iABIs, iAFs, and POLCs enhance pro-fibrotic and pro-inflammatory signaling pathways in the alveolar niche. (A) CellChat analysis indicates POLCs, KO POLCs, and iABIs are key players in the alveolar signaling niche of silica-treated (TR) Nkx2-1 KO (SKO) animals as they have among the strongest incoming and outgoing signals compared with silica-treated (TR) WT (SWT) animals. (B) Pro-fibrotic, pro-inflammatory, osteoclast-driving signaling pathways are enhanced in the SKO condition, including RANKL, SPP1, and TGFβ. (C) NicheNet analysis of the major ligand-receptor interactions between key players in the SKO niche (ABI, iABI, iAF, POLC, KO POLC) predicts that they signal reciprocally and engage in a pro-fibrotic loop that drives iABI and iAF emergence and immune cell recruitment. (D-G) Immunohistochemistry shows SIGLECF*+*/CD45*+* AMs are present in both injured and uninjured areas of TR lungs.

These predicted signaling interactions provided the third piece of evidence supporting an iABI, iAF, and POLC niche driving fibrosis in silica-treated Nkx2-1 KO mice. However, for these signaling interactions to be relevant, the predicted cellular partners need to be localized in common regions in the lung or within defined niches. iABI and iAF co-localized in fibrotic regions [**Fig.4I-J, Fig 5F-G]**, and lineaged labeld ABI/iABI and SiglecF^+^/CD45^+^ cells were found in close association in regions of fibrotic injury in silica treated Nkx2-1 KO lungs [**Fig.7D-G**]. Taken together, these data identify a physical and signaling niche driven by ABI cells which respond to inflammatory signals and activate pro-inflammatory and pro-fibrotic programs after lung injury [**Fig.8, Supplemental Figure 11]**. Since ABI are nearly absent from the WT lung after silica treatment, these data strongly support the conclusion that pre-existing, accumulated ABI participate in the lung injury response, potentiate tissue damage, and are therefore deleterious during lung injury and may impair tissue regeneration.

**Figure 8.**
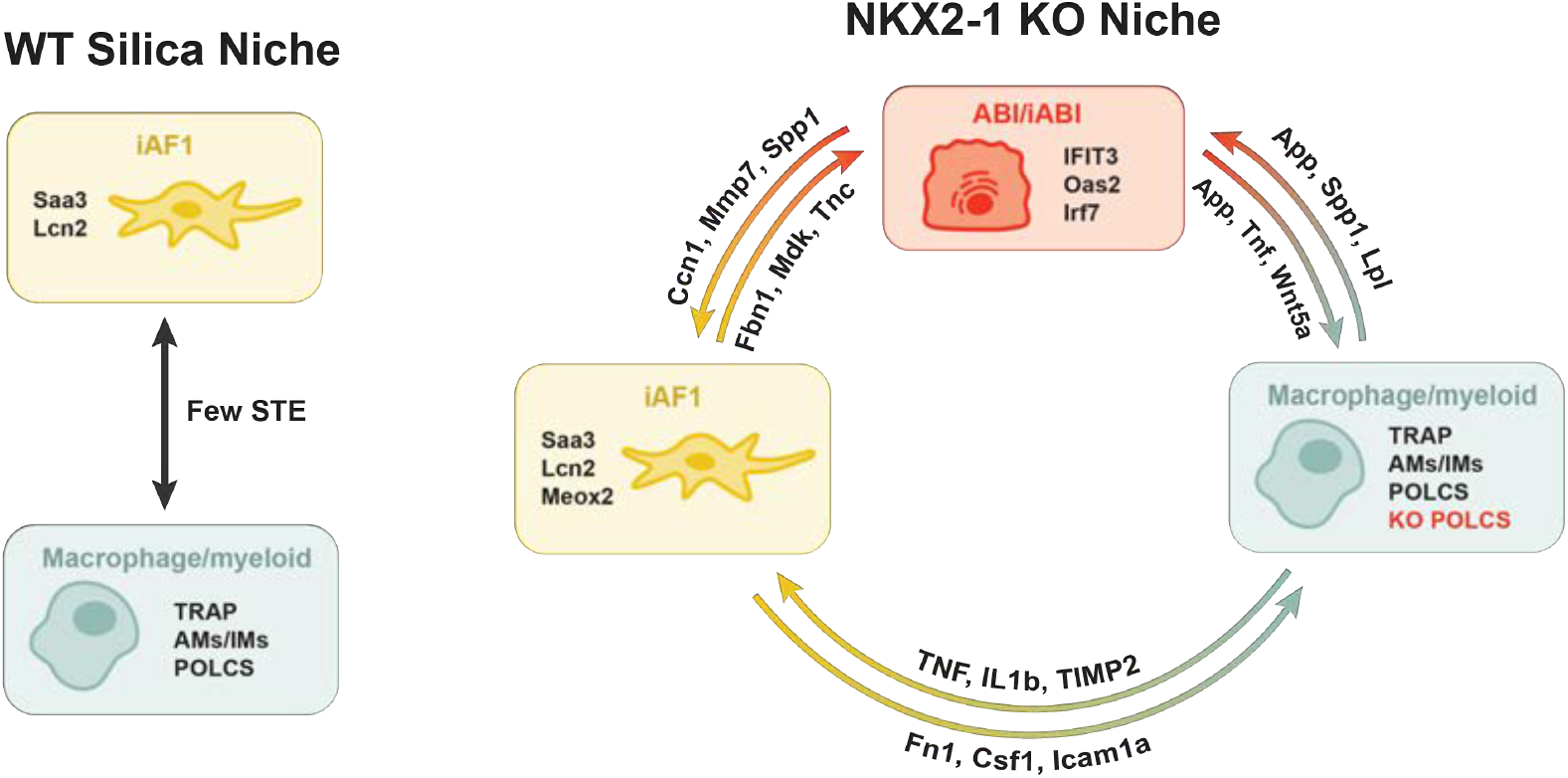
Accumulated ABI are activated by inflammation to drive lung fibrosis following silica exposure. In the WT niche, few ABI participate in a siganling loop between iAF and POLC cells to drive fibrotic nodule formation. In the Nkx2-1 niche, where ABI have accumulated, a fibrotic lung injury stimulus drives the emergence of inflammatory cell populations (iABI, iAF) that activate reciprocal ligand-receptor signaling interactions signaling between iABIs, iAFs, POLCs, and KO POLCS activates pro-fibrotic and pro-inflammatory signaling pathways in the alveolar niche.

## Discussion

The pathogenic versus protective nature of “transitional” cells has been a key unanswered and controversial question raised by recent studies. Here, we show that ABI cell accumulation in the lung is deletirious and drives a fibrotic lung injury milieu following induction of secondary injury, with increased lung damage, increased respiratory barrier permeability, and worsened fibrosis. Nkx2-1 deletion in AT2 progenitors enabled direct attribution of these effects to precisely timed accumulation of ABIs. While Nkx2-1 loss of function is an artificial method of ABI induction, reduction in Nkx2-1 expression and function is a common aspect of the transition of AT2 progenitors to Krt8^high^ cells in multiple models^10,13,27,57^. Our data demonstrate that Nkx2-1 null Krt8^high^ cells share a common molecular state with other reported ABI^25,58^, and these cells do not differentiate to other endoderm lineages^13^ or transition to more proximal lineages following injury. Instead, following secondary injury, Nkx2-1 null ABIs take on a uniquely inflamed state and directly interact with inflammatory alveolar fibroblasts and silica-specific macrophages to accumulate collagen and extracellular matrix proteins via stimulation of *Tgfb, Spp1, Collagen*, and *Laminin* pathways [**Fig.7,8**].

Present data position ABI accumulation as a key component of the pathogenesis of lung fibrosis^22^, and suggest that ABI represent a therapeutic target for future therapies. Our data imply that removal or rescue of ABI would prevent iABI fate transition, decreasing number and actvation of inflammatory alveolar fibroblasts, POLCs, and other pro-fibrotic cells. The cellular crosstalk between major inflammatory cell populations is critical to our proposed model in which ABIs promote fibrotic lung remodeling by increasing collagen and extracellular matrix deposition in the presence of ABI. Strongly supportive of this model is the recent report that ablation of Krt8^high^/Sprr1a^+^ ABI attenuates lung fibrosis in bleomycin-induced lung injury^59^. *Sprr1a* expression is associated with increase in Krt8^high^ cells during bleomycin injury, enabling specific ablation of these cells. However, not all Krt8^high^ cells express *Sprr1a*, and ablation with the diphtheria toxin/diphtheria-toxin receptor (DT/DTR) system effectively targeted only 40% of Sppr1a^+^ cells^59^. While the results of the Sprr1a cell ablation study support the conclusion that ABI are pathologic during lung fibrosis, its relevance must be considered in the context that bleomycin injury can cause spontaneously resolving lung fibrosis.

One key limitation of our model is the inability to clearly separate ABI accumulation from loss of AT2 progenitor function in the lung following secondary injury. We chose silica-induced fibrosis rather than bleomycin or other lung injury models to mitigate this limitation, since silica induces fibrosis primarily via activation of macrophage signaling and is associated with limited epithelial injury^30,31,35^. Present data demonstrate little accumulation of Krt8^high^ cells in WT animals treated with silica. These aspects improve the dynamic range of our conditions, since WT mice had minimal ABI accumulation and Nkx2-1 KO mice exhibited extensive and widespread accumulation. While beneficial for addressing the specific question of whether ABIs are deleterious, further studies are needed to identify key molecular partners of Nkx2-1 that prevent ABI accumulation or to target epigenetic or signaling pathways which could represent more tractable therapeutic targets than core transcriptional regulators. Our data emphasize the value of specific defiinition of cell states within heterogeneous Krt8^high^ cell populations, and suggest that future studies focused on a combination of inducible ABI gain and rescue would be required to enable the fine grained molecular phenotyping of ABI cell function that will be needed to move toward directed therapies to rescue ABI in patients with established ABI accumulation and fibrotic lung disease.

In the absence of ABI accumulation, the WT animals exposed to silica have much lower levels of iAFs and POLCs, emphasizing the role of the ABI cells in promoting these pathogenic cell states following injury. It is therefore tempting to speculate that ABI may also participate actively in the tissue response to other lung injuries, including viral pneumonia (SARS-COV2, influenza) and ARDS. Fibrotic responses can be beneficial in the setting of acute injury, helping to restore local tissue integrity and prevent progression of disease beyond the boundaries of the impacted region of the lung. The pro-fibrotic role of ABI may therefore be context dependent, beneficial in some injury/repair situations, and deleterious in others. While present data emphasize that ABI can be pathogenic, in the future, clear definition of the contexts in which ABI-induced fibrotic signaling is beneficial vs pathogenic will be required to identify patients who could most benefit from targeting of ABI for therapy in lung disease.

## Acknowledgements

The authors would like to thank the Single Cell Genomics Facility (especially Kelly Rangel and Shawn Smith), Bio-Imaging and Analysis Facility (especially director Matt Kofron), Genomics Sequencing Facility, and the Research Flow Cytometry Facility of the Cincinnati Children’s Research Foundation, all of whom have provided extensive technical support. We thank Jeffrey A. Whitsett for valuable editing of the manuscript.

## Author Contributions

Conceptualization – BZ, AT, WJZ

Data acquisition – BZ, HWN, KC, LP

Data analysis – BZ, HWN, KC, AE, ALZ, WJZ

Supervision – FXM, ALZ, WJZ

Writing – original draft – BZ, WJZ

Writing – review and editing - all authors

## Materials and Methods

Many of the following described methods correspond with previously published work in our laboratory^13,50^.

### Ethical compliance and animals

All animal studies were conducted following the guidelines of the Cincinnati Children’s Hospital Medical Center (CCHMC) Institutional Animal care and Use Committee (IACUC). Proposed studies were evaluated by the IACUC prior to performance. All studies were performed following CCHMC regulatory and biosafety protocols. The following mouse lines were used: Tfcp2l1^CreERT2^ (B6;129S-Tfcp2l1^tm1.1(cre/ERT2)Ovi^/J; Jackson Strain #028732), Nkx2-1^fl/fl^ (a gift from Shioko Kimura), and R26R^EYFP^ (B6.129×1-Gt(ROSA)26Sor^tm1(EYFP)Cos^/J; Jackson Strain #006148). For tamoxifen-induced Cre recombination, in mouse models, 6-8-week-old mice were given intraperitoneal (IP) injections of Tamoxifen (Sigma, T5648; dissolved in ethanol and resuspended in corn oil) at a dose of 50 mg/kg, twice every other day at the experimental time points previously mentioned.

### In vivo silica exposure

Silica particles (Sigma-Aldrich, St. Louis, MO, catalog number S5631; particle size: 80% between 1 and 5μm, 99% between 0.5 and 10μm) were boiled in 1 N HCl for 1 hour, washed with deionized H2O, and dried at 100°C. The particles were then heat sterilized at 200°C for 2 hours and suspended in sterile saline, as previously described^34^. The endotoxin content of the silica particles was silica (<1.0 pg/μg) as determined using the LAL Chromogenic Endotoxin Quantitation Kit (Thermo Fisher Scientific, Rockford, IL) according to the manufacturer’s instructions^34^. Prior to administration, 100mg of silica particles were resuspended in 1mL of 0.9% sodium chloride (saline) and subsequently vortexed and sonicated. During oropharyngeal (o.a.) silica administration, each mouse was first anesthetized with IP Ketamine/Xylazine solution (comprised of 9mg/mL Ketamine and 0.9mg/mL Xylazine) at a dose of 0.01mL/g and suspended by string on a procedure board at an approximately 60°C angle by the incisor teeth. The mouth was opened, the tongue was pulled forward, and 5mg of resuspended silica in saline was placed at the base of the tongue. The tongue was released after aspiration of the silica^34^.

### Mouse lung harvest

Mice were anesthetized via IP Ketamine/Xylazine solution, followed by euthanasia via cervical dislocation and thoracotomy. The chest cavity was opened to expose the heart and lungs. The right ventricle was perfused with 10mL of cold PBS to clear blood from the lungs. For flow cytometry, the right lobes were tied off, cut, and placed in PBS for downstream processing. For tissue fixation for histology and immunofluorescence, the trachea was cannulated, and lungs (or only the left lobe) were inflated via syringe using 2% paraformaldehyde (PFA). Inflated lung lobe(s) were immersed in either a conical or small glass jar of 2% PFA and left on a rocker at 4°C overnight^13,50^. For bronchoalveolar lavage fluid (BALF) collection, the trachea was cannulated, and the lungs were inflated with 1mL of BAL buffer (sterile, vacuum-filtered solution of DPBS, 2nM EDTA, and 0.5% of Fetal Bovine Serum^51^). The buffer was subsequently aspirated into a syringe, and this flushing procedure was repeated twice for a total of 2mL BALF collection per mouse.

### Processing fixed lung tissue for histology and immunofluorescence

Following inflation, lungs were trimmed of connective tissue and heart and place into histological cassettes. Tissue in cassettes were then washed 3X in DEPC-treated PBS, 1X in DEPC-treated 30% ethanol, 1X in DEPC-treated 50% ethanol, and 1X in DEPC-treated 70% ethanol. Using the standard CCHMC Pulmonary Biology automated processing protocol (Thermo Scientific, Excelsior ES), the samples were embedded in paraffin. Samples were sectioned at a thickness of 5μm with sections examined from the anterior to poster of lungs to ensure complete analysis of all regions of lung parenchyma. Paraffin sections were incubated at 65 °C for 2 h, deparaffinized in xylene (3x for 10 min), rehydrated through an ethanol gradient, and examined following standard H&E staining. Slides were mounted with Permount Mounting Medium (Electron Microscopy Sciences, 17986-05) and cover-slipped with #1.5 Gold Seal 3419 Cover Glass (Electron Microscopy Sciences, 63790-01). For pentachrome staining, Movat Pentachrome Stain Kit (Abcam, ab245884) was used according to manufacturer’s instructions. Immunofluorescence on paraffin sections was performed as previously described^9,13^. Slides underwent deparaffinization, were rehydrated, and sodium citrate antigen retrieval was performed to prepare for staining (10 mM, pH 6.0), and blocking. Immunofluorescence was performed on paraffin sections using antibodies in Supplementary Table 2 and the following reagents: ImmPRESS® HRP Horse Anti-Rabbit IgG Polymer Detection Kit (Vector Labs, MP-7401-50) or ImmPRESS® HRP Goat Anti-Rat IgG Polymer Detection Kit (Vector Labs, MP-7404-50). Following the application of TSA fluorophores (listed in Supplementary Table 2; 1:100), sections were stained with DAPI (Invitrogen, D1306; 1:1000) and mounted using Prolong Gold antifade mounting medium (Invitrogen, P36930).

### Alveolar epithelial permeability tests

Bronchoalveolar lavage fluid (BALF) was diluted 1:250,000, and albumin concentrations were measured across 2 technical replicates with the Mouse Albumin ELISA Kit (Fortis Life Sciences, E99-134). BALF was diluted 1:10, and IgM concentrations were measured across 2 technical replicates with the Mouse IgM ELISA Kit (Fortis Life Sciences, E99-101). For both assays, absorbance measurements were read at 450nm, and concentrations were calculated accordingly from a standard curve.

### Hydroxyproline measurements of the murine lung

The right accessory lobe was flash frozen during mouse lung harvest and thawed prior to assay use for 15min on ice. Samples were homogenized in 100uL molecular-grade H2O and subsequently mixed with 100uL 12 M HCl. Samples were hydrolyzed at 120°C for 3 hours and centrifuged at 10,000g for 3 min. Hydroxyproline content was measured across 2 technical replicates per sample from the supernatant using the Hydroxyproline Assay Kit (Sigma-Aldrich, MAK-569-1KT) according to the manufacturer’s instructions.

### Lung damage assessment

Tile scans of H&E-stained left lobes were processed in ImageJ to ensure all JPEG images were of the same size and scale. Processed images were then run through an RStudio program as previously described, and “normal,” “damaged,” and “severe” surface area percentages were distinguished by pixel density determinations of the program^35^.

### Mouse lung digestion and single-cell suspension

Single cell suspensions from murine lungs were generated as previously described13. After euthanasia via IP Ketamine/Xylazine, the thorax was exposed, and the right ventricle of the heart was perfused with 10 mL of cold 1X PBS. All lung lobes were cut and submerged in ice-cold PBS. Lung tissue was minced and transferred to a gentleMACS C Tube (Miltenyi 130-096-334) before 5 mL of 37°C digest buffer mixture (dispase (Corning 354235), DNase1 (GoldBio D-301), collagenase type 1 (Gibco 17100017), and 1X PBS) was added. C tubes were placed on a gentleMACS Octo Dissociator (Miltenyi Biotec, 130-096-427) and the protocols were run: “m_lung_01_02” (36 s) X2, “37C_m_LIDK_1” (36 min 12 s) X1, and “m_lung_01_02” (36 s) X1. All samples were then passed through a 70uM filter and centrifuged at 800g for 8 minutes at 4°C. The supernatant was aspirated, and remaining cells were incubated in room temperature RBC Lysis Buffer (eBioScience 00-4333-57) for 5 minutes. Samples were centrifuged at 500g for 5 minutes at 4°C, the supernatant was aspirated, and the cell pellet was resuspended in cold MACS Buffer [AutoMACS Rinsing Buffer (Miltenyi 130-091-222) + MACS BSA Stock Solution (Miltenyi 130-091-376)] before being passed through a 40uM filter and washed with 5 mL of MACS Buffer. Samples were centrifuged at 500g for 5 minutes at 4°C and resuspended in a final volume of 5 mL of MACS buffer to generate a single cell suspension.

### Fluorescence-activated cell sorting (FACS) preparation and processing

Cells were incubated in Fc Receptor Binding Inhibitor Polyclonal Antibody (eBioScience 14-9161-73) diluted 1:100 in MACS Buffer for 10 minutes at room temperature. After incubation, samples were centrifuged, supernatant was removed, and cells were incubated in CD326 APC antibody (Invitrogen 17-5791-82) diluted 1:100 in MACS Buffer at room temperature for 10 minutes in the dark. After another round of centrifuging and aspirating, cells were washed with 5 mL of cold MACS Buffer, then centrifuged again. The supernatant was removed, and cells were incubated with Fixable Viability Dye eFluor™ 780 (eBioScience 65-0865-14) diluted 1:1000 in MACS Buffer for 15 minutes at room temperature in the dark. Cells were then centrifuged and washed 2-3X with 5 mL of cold MACS Buffer before being resuspended in a cell count adjusted volume of MACS Buffer and strained through 35uM filter lids of polystyrene FACS tubes on ice (Corning 352235) for sorting.

Using single-stain compensation beads (Invitrogen 01111142), gating was adjusted to remove debris and doublets using a BD FACSAria Fusion or BD FACSymphony cell sorter fitted with a 100uM nozzle. Live/CD326+/eYFP+ (AEP) cells were sorted into a 1.5 mL tube pre-coated in 500μL of cold SAGM (Lonza CC-3118) media to preserve cell viability. This protocol yields approximately 2x105 AT2 cells per mouse.

### Spectral flow cytometric analysis

Lungs were digested, and single-cell suspensions were generated as described. Cells were incubated in Mouse BD Fc Block (BD Pharmingen, 553141) at 1:100 for 10min at room temperature (RT). Cells were then washed with MACS buffer, centrifuged, and resuspended in antibody panel listed in Supplementary Table 2. Cells were washed, centrifuged, and strained through 35uM filter lids of polystyrene FACS tubes. Samples were then run on a 5-laser Cytek Aurora with unstained and single-color controls for spectral unmixing and compensation. Unmixing was done using Cytek’s SpectroFlo software; compensation was adjusted, and gating and analysis was performed using FlowJo.

### Sequencing/library preparation

From each single-cell suspension described above, a maximum of 16,000 cells were loaded into a channel of a 10x Genomics Chromium system by the Cincinnati Children’s Hospital Medical Center Single Cell Sequencing Core. Libraries for RNA (v3) and 10x Chromium Single Cell Gene Expression Flex were generated following the manufacturer’s protocol. Sequencing was performed by the Cincinnati Children’s Hospital DNA Sequencing Core using Illumina reagents. Raw Sequencing data was aligned to the mouse reference genome mm10 with CellRanger 3.0.2 to generate expression count matrix files. To detect YFP expressing cells following Cre-mediated activation, a YFP contig was added to the mm10 genome following 10x Genomics “Build a Custom Reference” instructions with modifications. Briefly, a custom EYFP.fasta file was generated using the ‘EYFP’ segment (682-1389) of the pEYFP-N1 plasmid sequence available through Addgene. This sequence was integrated into the standard mm10 assembly available from Ensembl to create a reference compatible for alignment with the CellRanger pipeline described above^13^.

### scRNA seq analysis and visualization

For scRNA-seq analysis, the output data from CellRanger was partitioned by Velocyto into spliced and unspliced reads. Spliced transcripts were used as the expression input in Seurat. We excluded cells with <2000 or >8000 features, and cells were clustered via the standard Seurat workflow. DoubletFinder was used to identify and remove putative doublets. We integrated the libraries from individual time points and treatments in Seurat, then reclustered, generated a UMAP project, and identified samples based on expression similarity^13^.

### Flex analysis and visualization

After Harmony normalization and initial clustering with Seurat v5^52^, preliminary cluster identifications were made based on marker gene expression. Based on these preliminary identifications, cells were assigned to one of 5 major groups: epithelium, endothelium, mesenchyme, monocytes, leukocytes, and neutrophils/basophils, to allow for better resolution. Cells in these groups were re-clustered and assigned final identities based on marker gene expression and treatment condition of the clusters. Marker genes for novel clusters, and enriched (fold-change >2, p<0.005) or depleted genes between clusters of interest were identified using Seurat. Using these enriched or depleted genes, Metascape analysis^53^ was used to identify GO terms, potentially relevant protein-protein interactions, and transcription factors with enriched motifs upstream of changed genes. CellChat^47,48^ and NicheNet^41^ were used to identify interactions between all clusters and inflammatory clusters of interest, respectively.

## SUPPLEMENTARY MATERIALS

**Supplementary Table 4-1.**
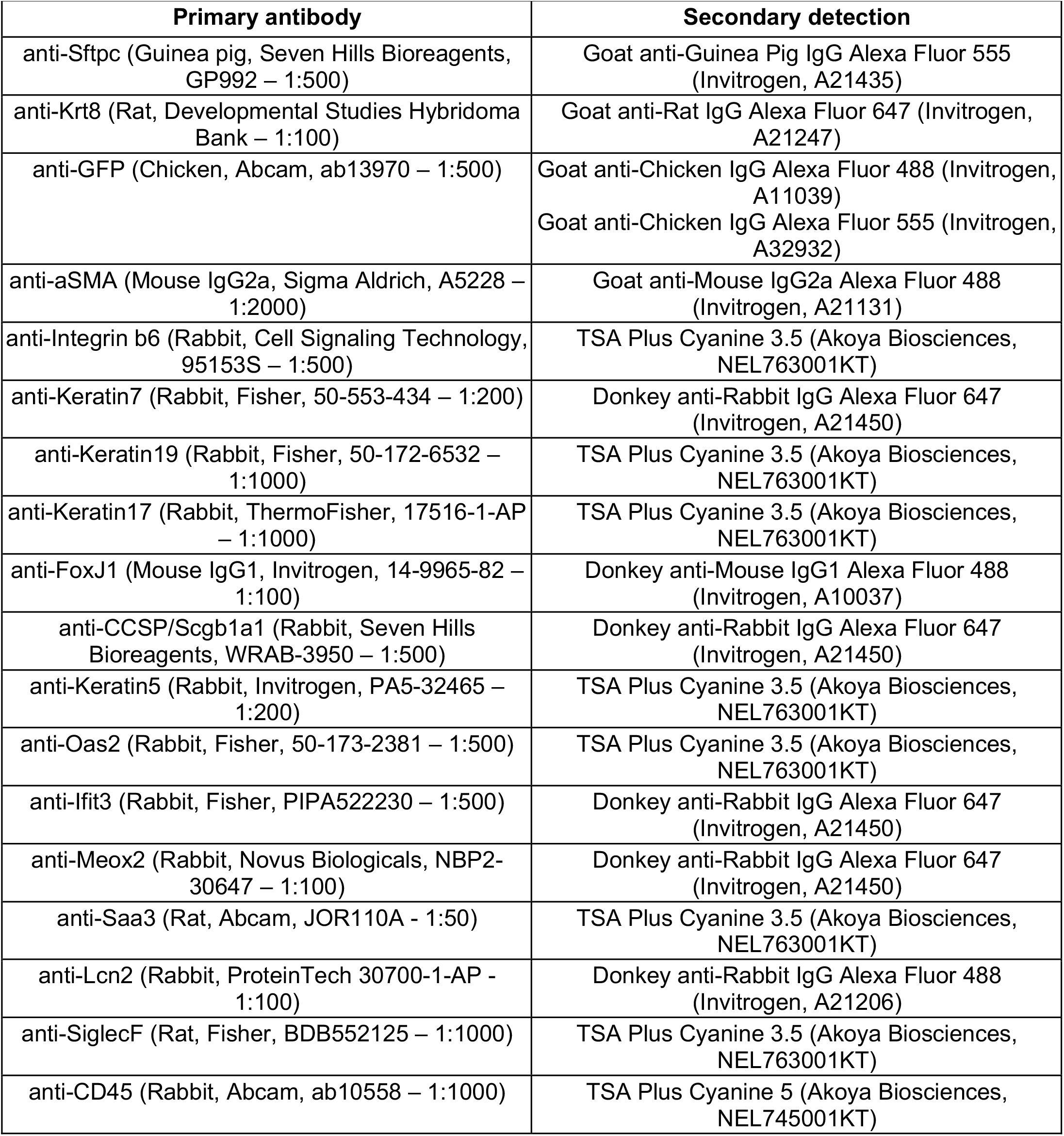
Antibodies for immunofluorescence on paraffin sections.

| Primary antibody | Secondary detection |
| --- | --- |
| anti-Sftpc (Guinea pig, Seven Hills Bioreagents, GP992 – 1:500) | Goat anti-Guinea Pig IgG Alexa Fluor 555 (Invitrogen, A21435) |
| anti-Krt8 (Rat, Developmental Studies Hybridoma Bank – 1:100) | Goat anti-Rat IgG Alexa Fluor 647 (Invitrogen, A21247) |
| anti-GFP (Chicken, Abcam, ab13970 – 1:500) | Goat anti-Chicken IgG Alexa Fluor 488 (Invitrogen, A11039)<br>Goat anti-Chicken IgG Alexa Fluor 555 (Invitrogen, A32932) |
| anti-aSMA (Mouse IgG2a, Sigma Aldrich, A5228 – 1:2000) | Goat anti-Mouse IgG2a Alexa Fluor 488 (Invitrogen, A21131) |
| anti-Integrin b6 (Rabbit, Cell Signaling Technology, 95153S – 1:500) | TSA Plus Cyanine 3.5 (Akoya Biosciences, NEL763001KT) |
| anti-Keratin7 (Rabbit, Fisher, 50-553-434 – 1:200) | Donkey anti-Rabbit IgG Alexa Fluor 647 (Invitrogen, A21450) |
| anti-Keratin19 (Rabbit, Fisher, 50-172-6532 – 1:1000) | TSA Plus Cyanine 3.5 (Akoya Biosciences, NEL763001KT) |
| anti-Keratin17 (Rabbit, ThermoFisher, 17516-1-AP – 1:1000) | TSA Plus Cyanine 3.5 (Akoya Biosciences, NEL763001KT) |
| anti-FoxJ1 (Mouse IgG1, Invitrogen, 14-9965-82 – 1:100) | Donkey anti-Mouse IgG1 Alexa Fluor 488 (Invitrogen, A10037) |
| anti-CCSP/Scgb1a1 (Rabbit, Seven Hills Bioreagents, WRAB-3950 – 1:500) | Donkey anti-Rabbit IgG Alexa Fluor 647 (Invitrogen, A21450) |
| anti-Keratin5 (Rabbit, Invitrogen, PA5-32465 – 1:200) | TSA Plus Cyanine 3.5 (Akoya Biosciences, NEL763001KT) |
| anti-Oas2 (Rabbit, Fisher, 50-173-2381 – 1:500) | TSA Plus Cyanine 3.5 (Akoya Biosciences, NEL763001KT) |
| anti-Ifit3 (Rabbit, Fisher, PIPA522230 – 1:500) | Donkey anti-Rabbit IgG Alexa Fluor 647 (Invitrogen, A21450) |
| anti-Meox2 (Rabbit, Novus Biologicals, NBP2-30647 – 1:100) | Donkey anti-Rabbit IgG Alexa Fluor 647 (Invitrogen, A21450) |
| anti-Saa3 (Rat, Abcam, JOR110A - 1:50) | TSA Plus Cyanine 3.5 (Akoya Biosciences, NEL763001KT) |
| anti-Lcn2 (Rabbit, ProteinTech 30700-1-AP - 1:100) | Donkey anti-Rabbit IgG Alexa Fluor 488 (Invitrogen, A21206) |
| anti-SiglecF (Rat, Fisher, BDB552125 – 1:1000) | TSA Plus Cyanine 3.5 (Akoya Biosciences, NEL763001KT) |
| anti-CD45 (Rabbit, Abcam, ab10558 – 1:1000) | TSA Plus Cyanine 5 (Akoya Biosciences, NEL745001KT) |

**Supplementary Table 4-2.** Spectral flow cytometry antibody panel.

| Antibody | Supplier | Dilution |
| --- | --- | --- |
| Podoplanin-BV421 (8.1.1) | Biolegend, 127423 | 1:100 |
| CD31-eFluor450 (390) | ThermoFisher (Invitrogen), 48-0311-82 | 1:100 |
| MHCII-BV480 (M5/114.15.2) | Fisher (BD Horizon), BDB566088 | 1:300 |
| CD326-BV785 (G8.8) | Biolegend, 118245 | 1:100 |
| CD24-PerCP/Cy5.5 (M1/69) | Biolegend, 101823 | 1:100 |
| CD200-PE/Dazzle594 (OX2) | Biolegend, 123819 | 1:300 |
| CD326-APC (G8.8) | ThermoFisher (Invitrogen), 17-5791-82 | 1:100 |
| ItgB4-AlexaFluor647 (346-11A) | Biolegend, 123607 | 1:100 |
| CD45-AlexaFluor700 (30-F11) | Biolegend, 103127 | 1:200 |
| SiglecF APC/Cy7(E50-2440) | Fisher (BD Biosciences), BDB565527 | 1:200 |
| CD11c PE (N418) | BioLegend, 117305 | 1:400 |
| CD11b BV510 (M1/70) | BioLegend, 101245 | 1:200 |
| CD64/FcyRI APC (S18017D) | BioLegend, 1611005 | 1:100 |
| Ly6C APC/Fire 810 (HK1.4) | BioLegend, 128055 | 1:400 |
| Ly6G Spark UV 387 (1A8) | BioLegend, 127677 | 1:100 |

**Supplementary Figure 1.**
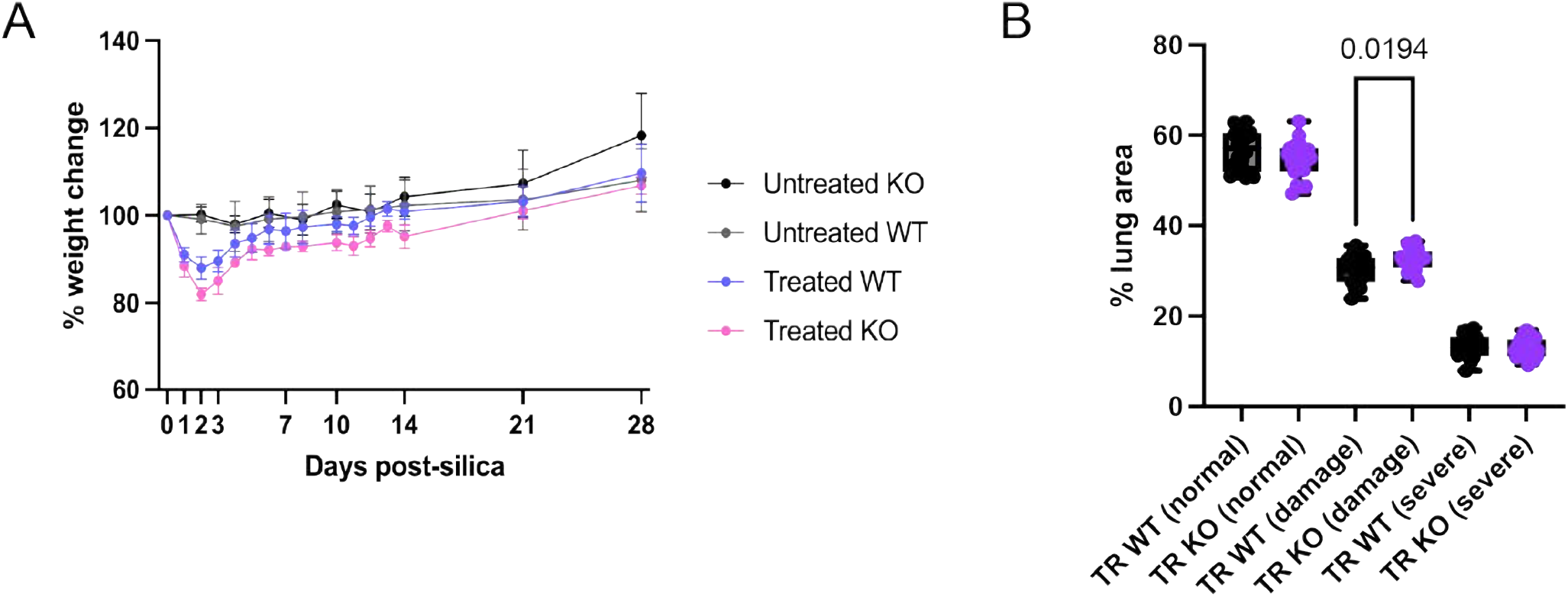
(A) Percentage weight change between untreated and silica-treated conditions. (B) Percentage of lung area characterized as “normal,” “damaged,” and “severe” across all collected samples. Exact p-values calculated via one-way ANOVA with multiple comparisons.

**Supplementary Figure 2.**
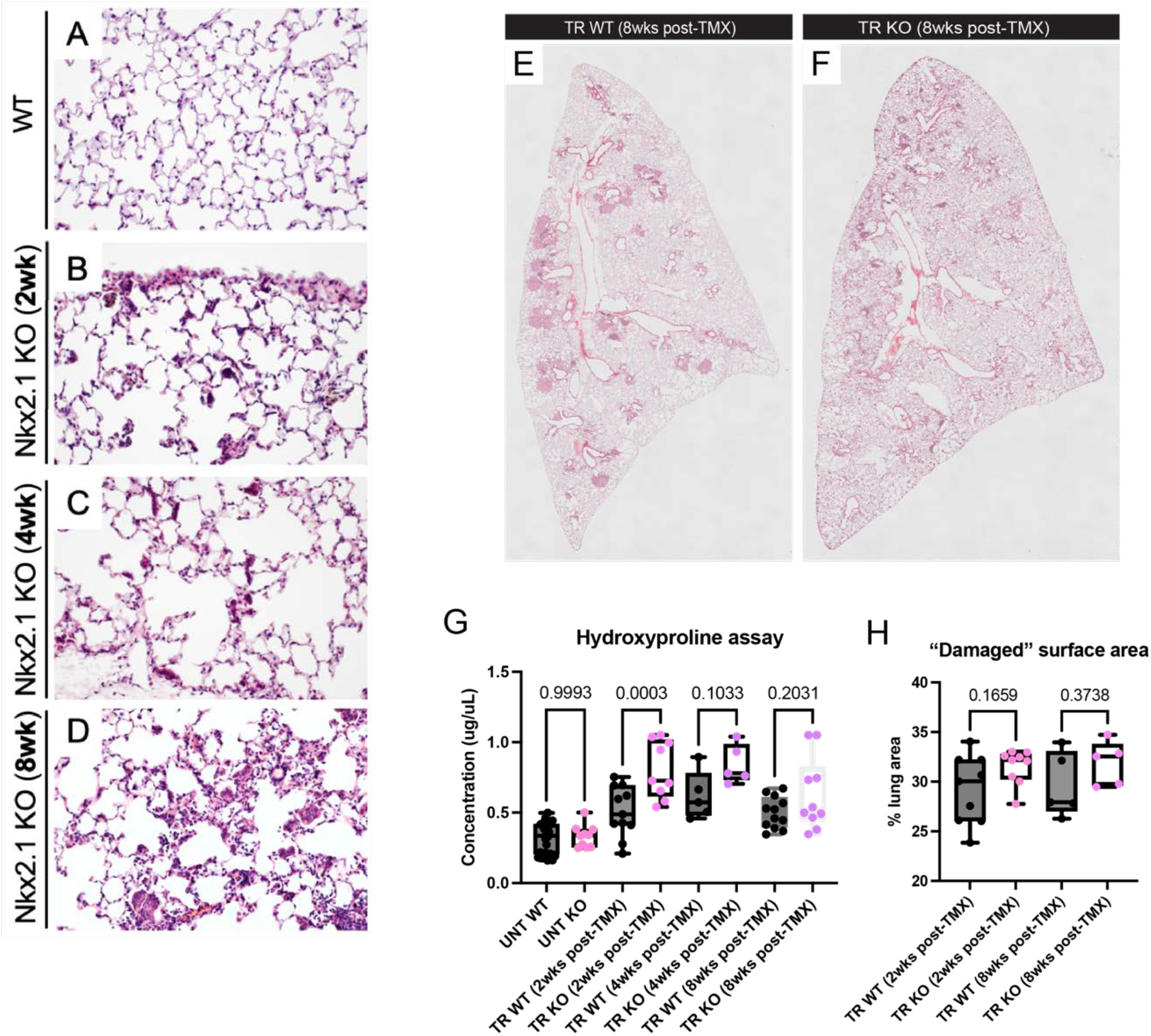
(A-D) H&Es exhibit progressively enlarged multi-cellular Krt8^high^ cell clusters 2weeks-(B), 4weeks- (C), and 8weeks- (D) post-tamoxifen-induced Cre recombination of Nkx2-1 floxed alleles in AEPs. (E,F) Silica-treated WT and silica-treated KO lungs 8wks post-TMX-induced Cre recombination present similarly to phenotype described in main Figure 2, which shows TR WT and TR KO lungs 2wks post-TMX-induced Cre recombination. (G) Hydroxyproline assay demonstrates upregulated collagen deposition in TR KO lungs at all treated time points after TMX-induction, with the most statistically significant upregulation being 2wks post-TMX-induced Cre recombination. (H) “Damaged” surface area quantified by separating samples based on time points demonstrates common trend of increased percentage of “damaged” surface area in TR KO lungs. All exact p-values calculated via one-way ANOVA with multiple comparisons.

**Supplementary Figure 3.**
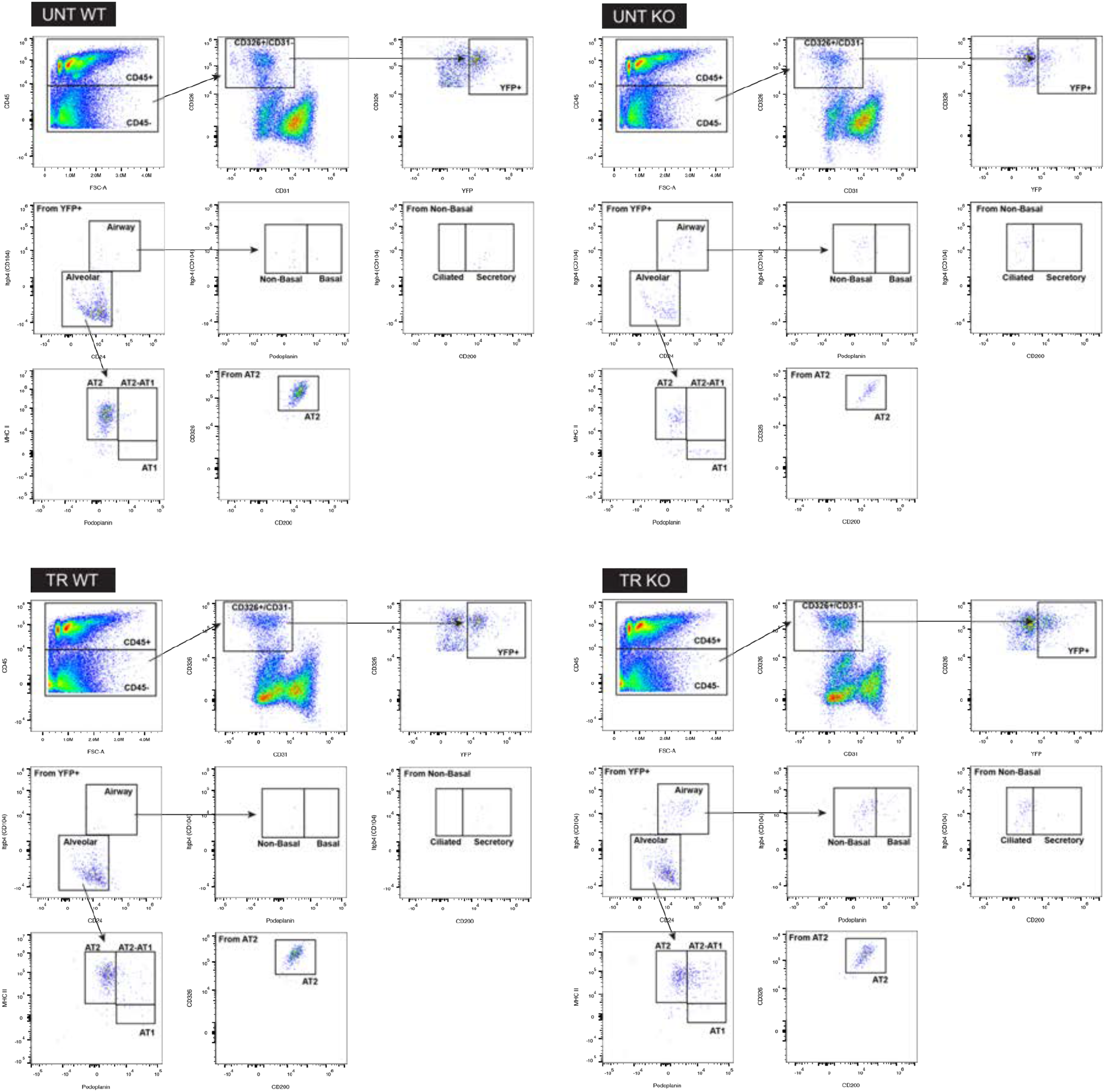
Spectral flow cytometry gating of alveolar and airway epithelial cells with representative gating from each experimental condition using panel described in Supplemental Table 2.

**Supplementary Figure 4.**
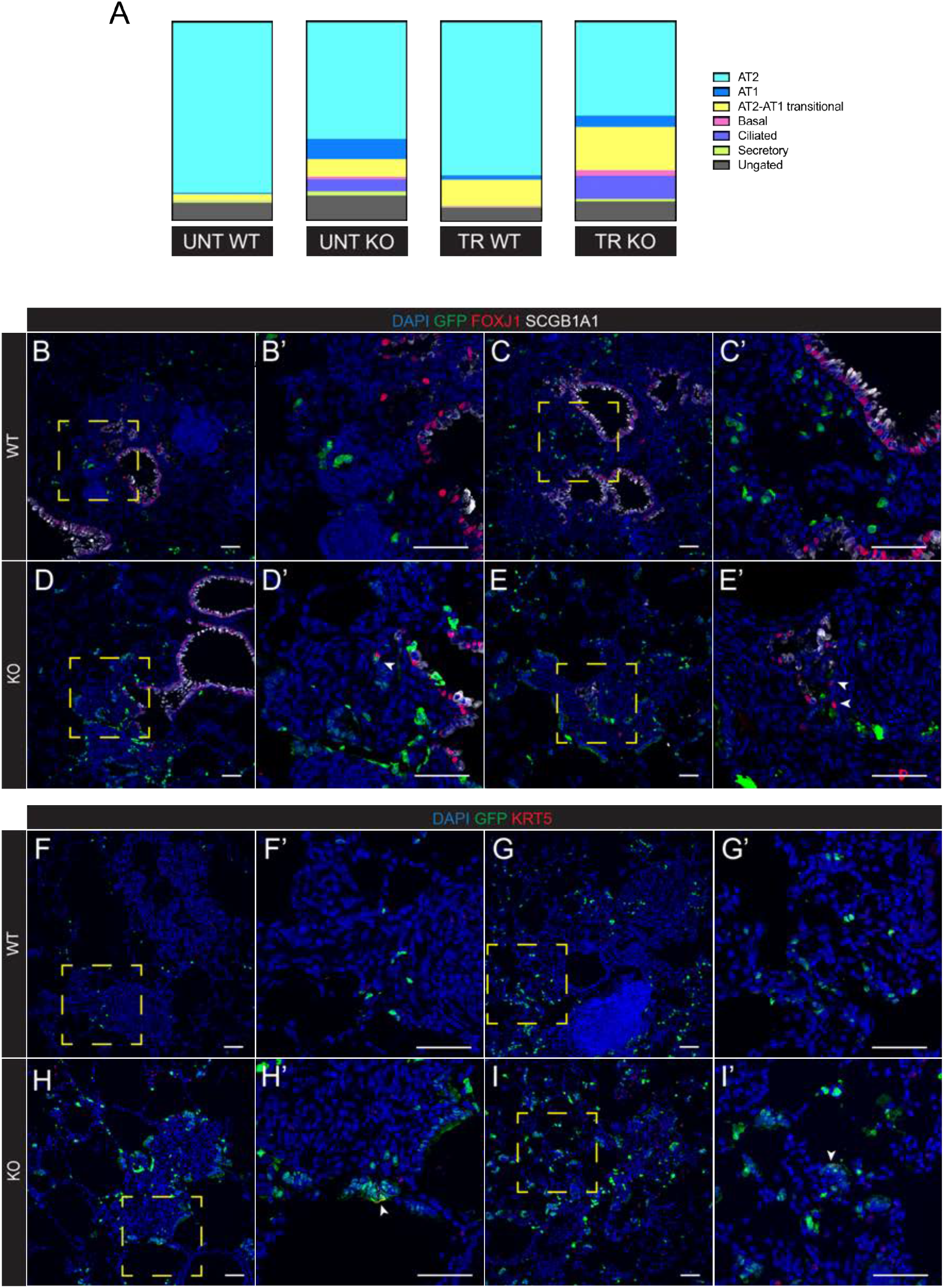
(A) Spectral flow cytometry characterization of airway epithelial cell fate in Nkx2-1 KO conditions. These data predicted significant transition to ciliated cells and basal cells. (B-E) Immunohistochemistry demonstrates that very few lineage-labeled Krt8^high^ cells express ciliated cell marker FOXJ1 in TR KO condition (arrows). (F-I) Immunohistochemistry also demonstrates that very few lineage-labeled Krt8^high^ cells express basal cell marker KRT5 in TR KO condition (arrows). We conclude that the potential ciliated and basal cells are likely ABI. Scale bars = 50 micron.

**Supplementary Figure 5.**
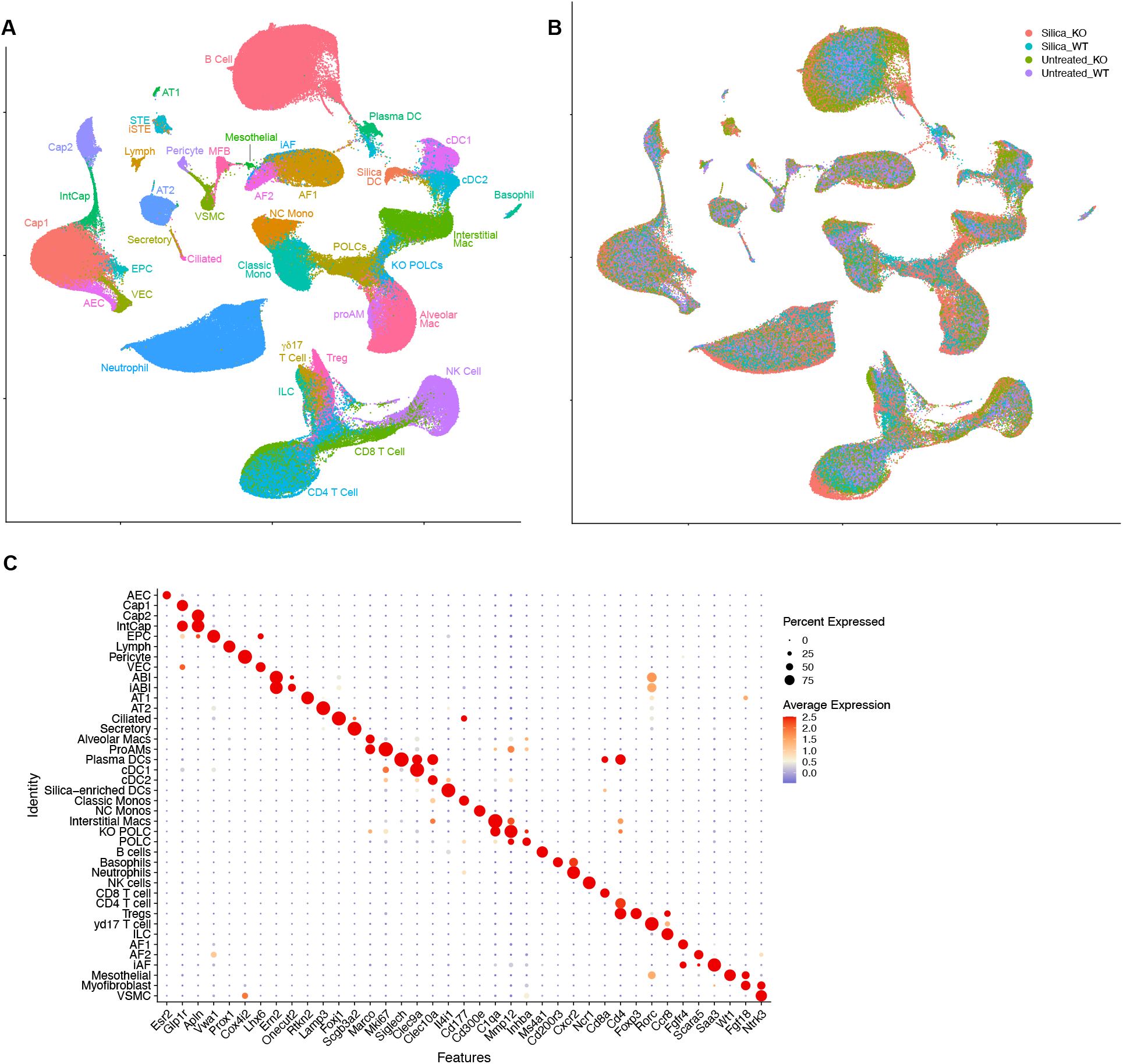
A) UMAP of integrated analysis of all identified cell populations concatenated across whole lung scRNA sequencing of all experimental conditions. B) UMAP as in A, but colored by experimental condition. C) Dot plot of highly enriched genes for each identified cell type from cells shown in A.

**Supplementary Figure 6.**
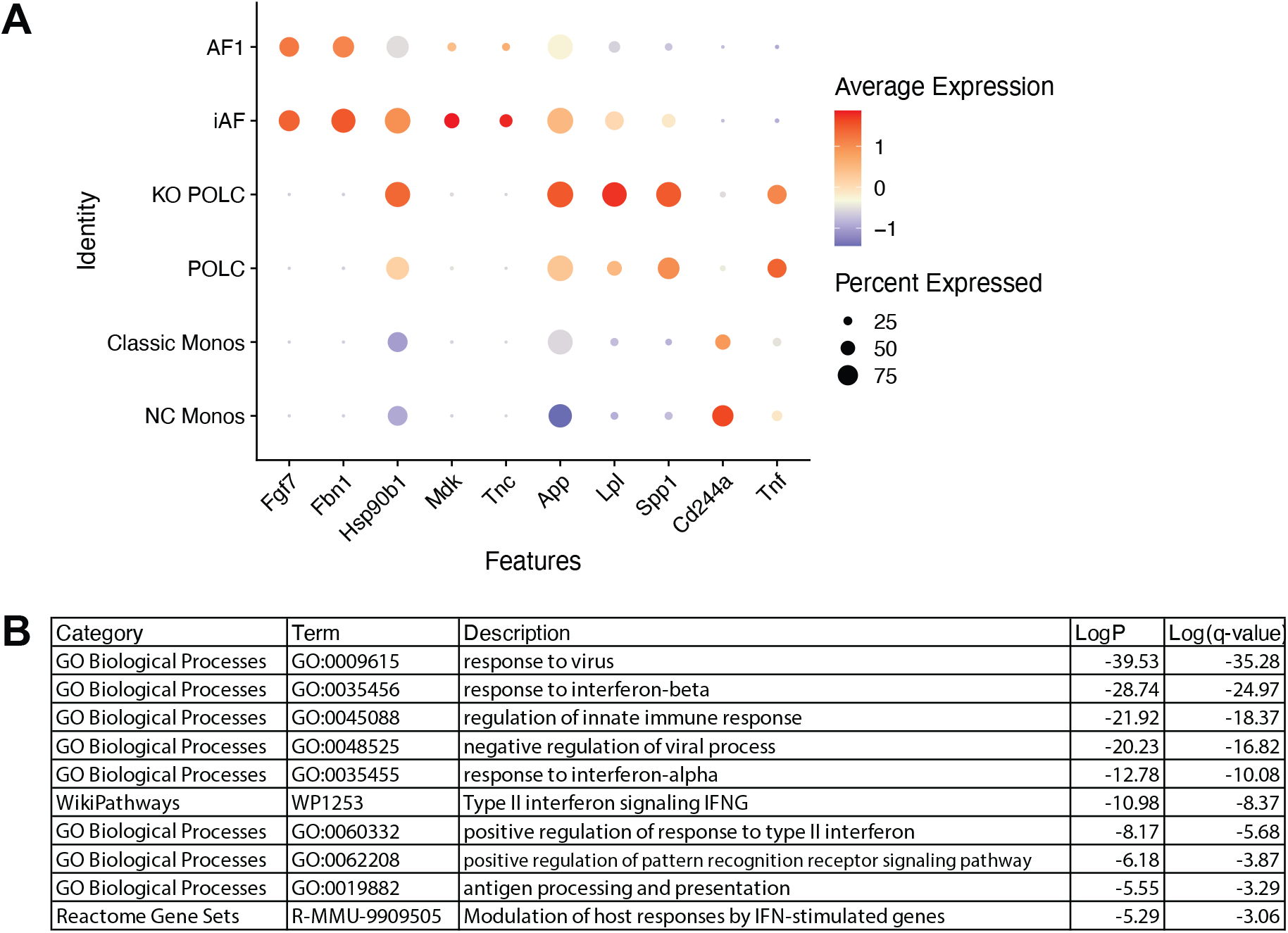
A) Dot plot showing relative enrichment of ligands predicted to trigger the iABI cell state in the cells that express these ligands as identified by NicheNet analysis (Figure 4F). Color shows the relative enrichment by cell type (red = enriched, blue = depleted) while the size of the circle indicates the fraction of cells in a cluster that express the gene of interest. B) Gene Ontology (GO) and Reactome terms and pathways enriched in iABI relative to ABI cells based on a Metascape analysis of significantly enriched genes up at least two-fold.

**Supplementary Figure 7.**
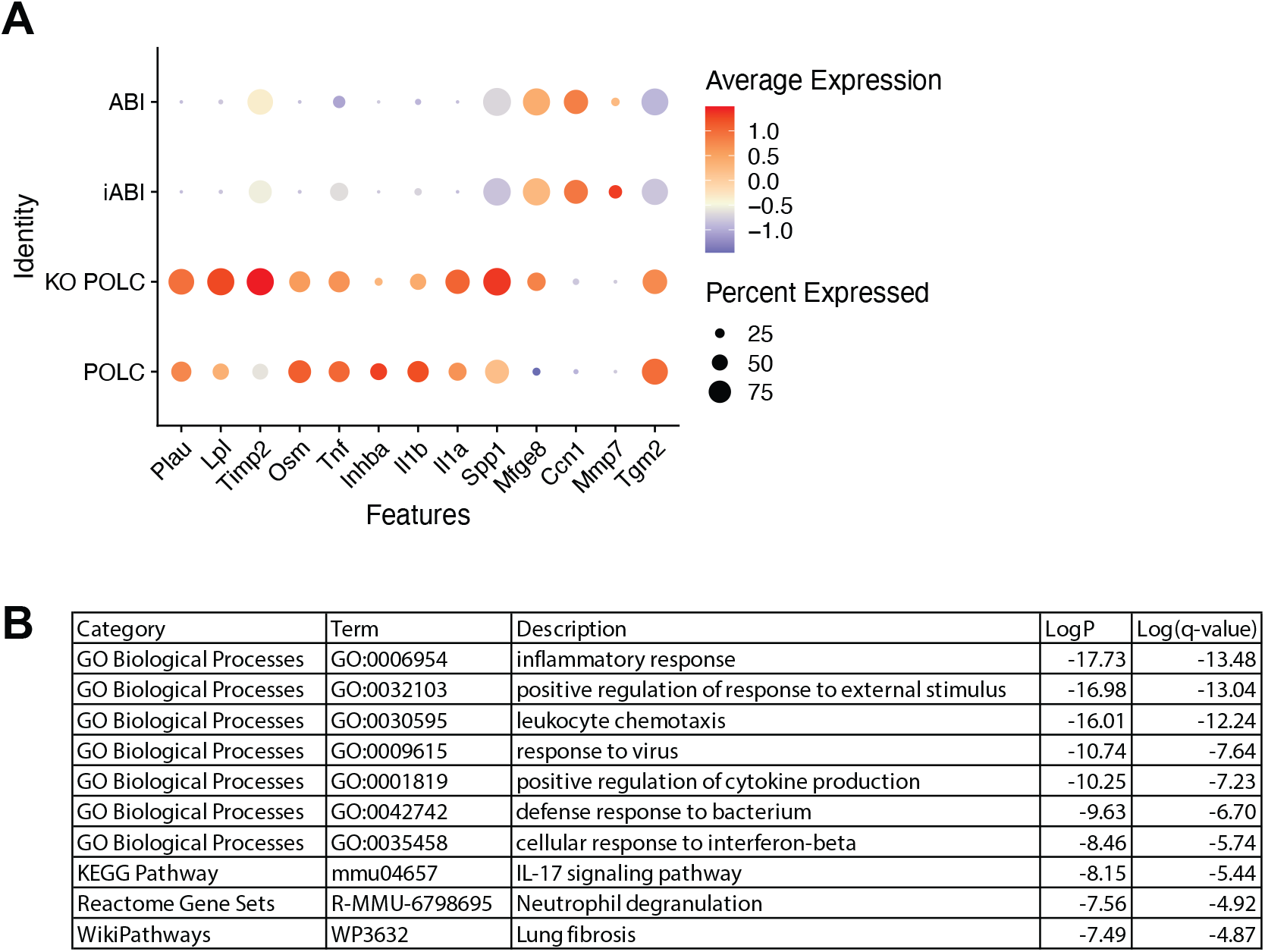
A) Dot plot showing relative enrichment of ligands predicted to trigger the iAF cell state in the cells that express these ligands as identified by NicheNet analysis (Figure 5I). Color shows the relative enrichment by cell type (red = enriched, blue = depleted) while the size of the circle indicates the fraction of cells in a cluster that express the gene of interest. B) Gene Ontology (GO) and Reactome terms and pathways enriched in iAF relative to AF1 cells based on a Metascape analysis of significantly enriched genes up at least two-fold.

**Supplementary Figure 8.**
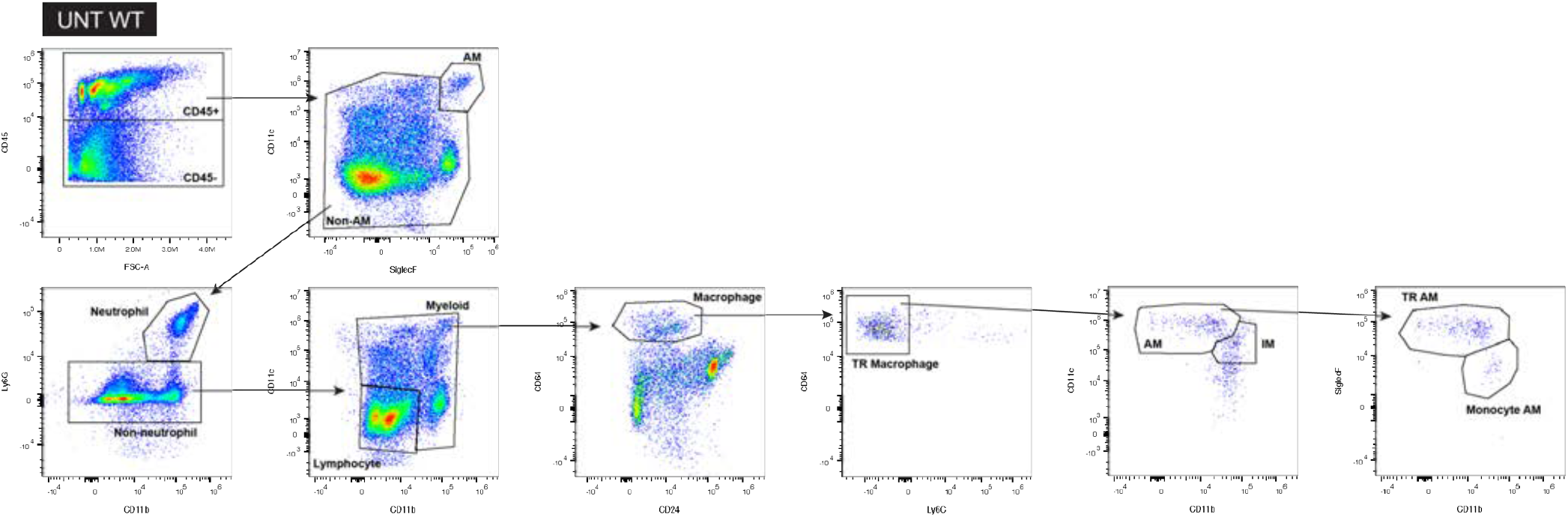
Spectral flow cytometry gating from representative untreated (UNT) WT sample showing delineation of alveolar macrophages (AM), tissue-resident AMs, monocyte-derived AMs, and tissue-resident interstitial macrophages (IM). Immune cell flow cytometry gating is shown as previously described^54^.

**Supplementary Figure 9.**
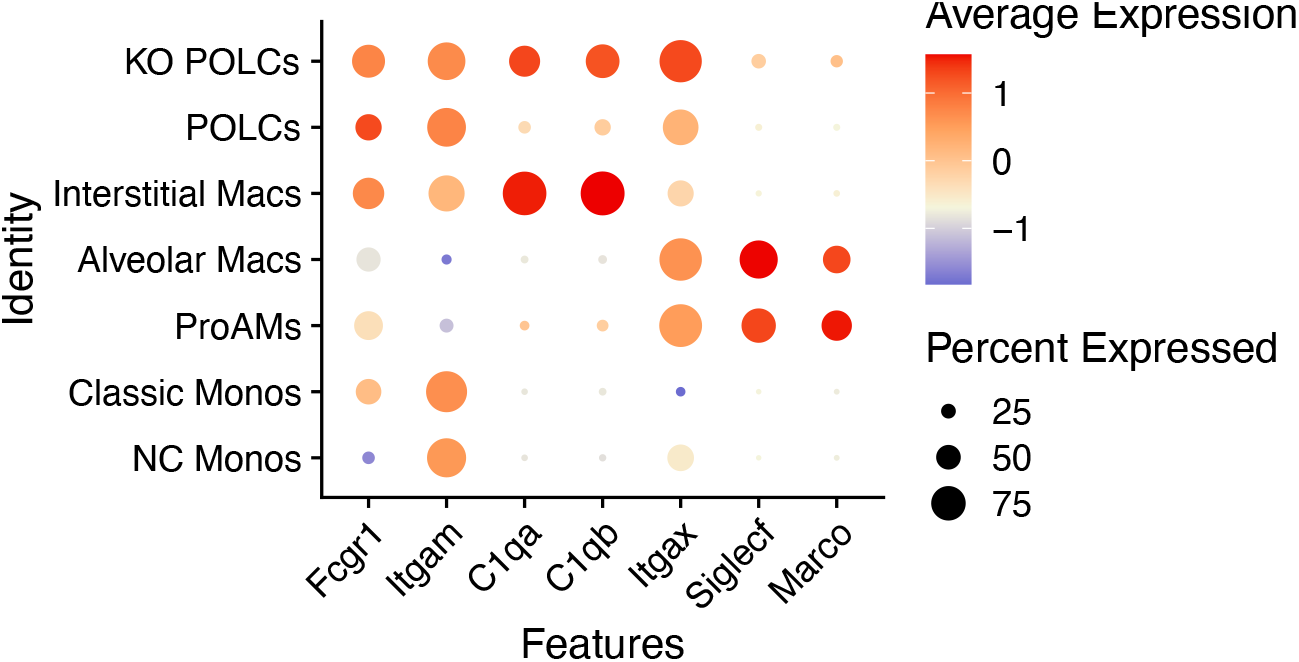
Dot Plot comparing marker gene expression between POLCs, KO POLCs, and other key immune cell populations.

**Supplementary Figure 10.**
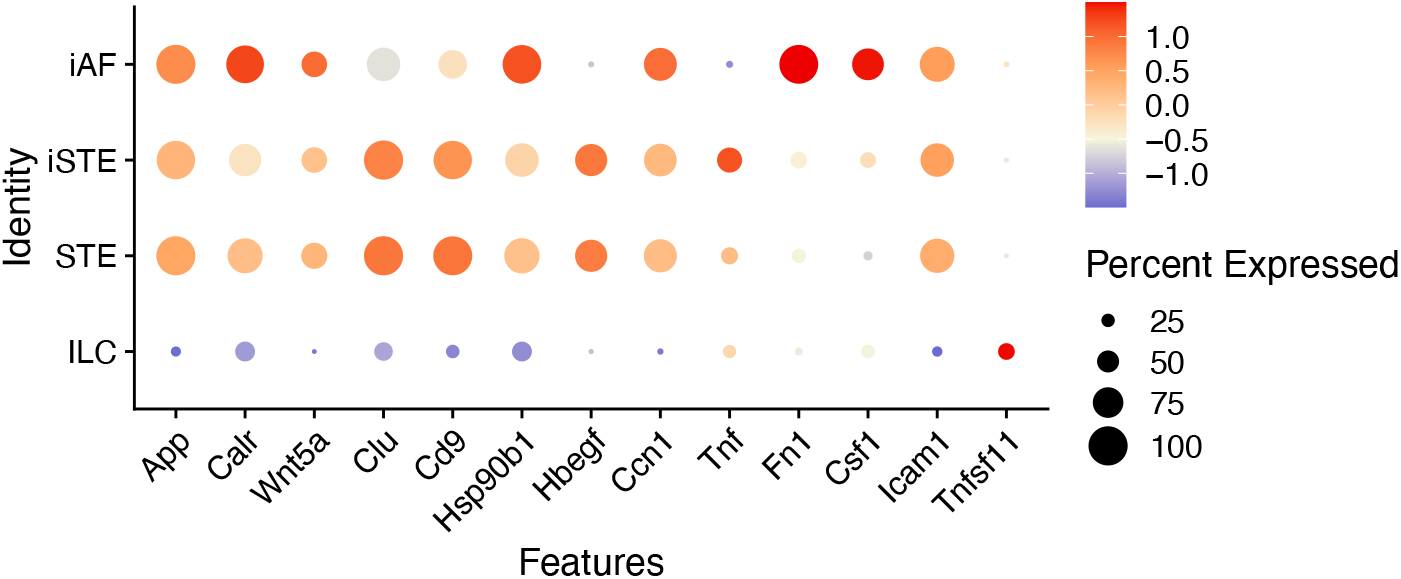
A) Dot plot showing relative enrichment of ligands present in the KO niche as identified by NicheNet analysis (Figure 7C). Color shows the relative enrichment by cell type (red = enriched, blue = depleted) while the size of the circle indicates the fraction of cells in a cluster that express the gene of interest. ILC = innate lymphoid cells

**Supplementary Figure 11.**
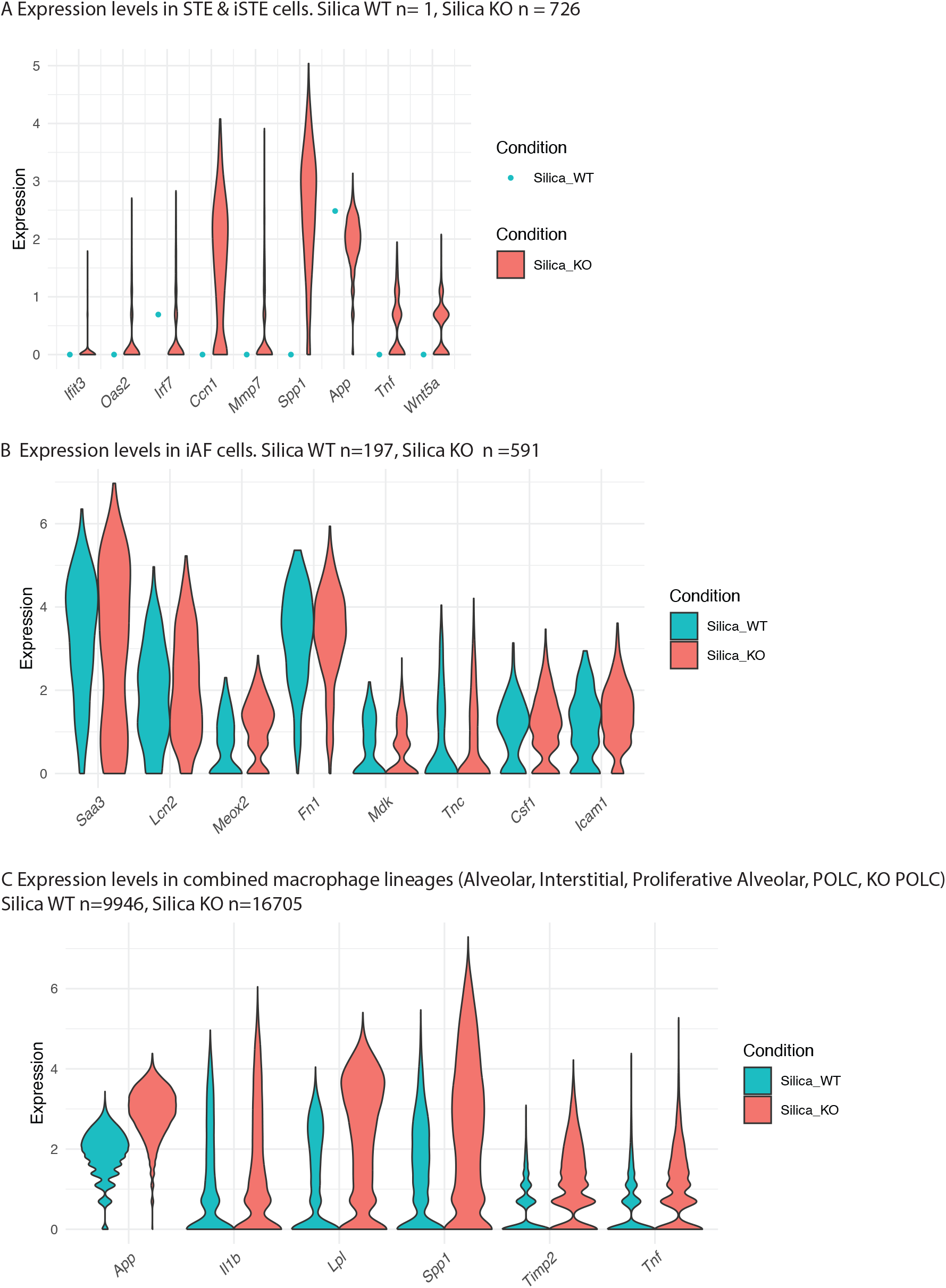
Violin plots showing the relative gene expression levels for genes in Figure 8, split by genotype and cell type. A) iSTE and STE, B) iAF, C) combined macrophage lineages. Number of cells identified of each type per condition noted.

